# Signaling Node at TSC2 S1365 Potently Regulates T-Cell Differentiation and Improves Adoptive Cellular Cancer Therapy

**DOI:** 10.1101/2022.11.30.518569

**Authors:** Chirag H Patel, Yi Dong, Brittany L Dunkerly-Eyring, Jiayu Wen, Mark J Ranek, Laura M Bartle, Daniel B Henderson, Jason Sagert, David A Kass, Jonathan D Powell

## Abstract

MTORC1 integrates signaling from the immune microenvironment to regulate T-cell activation, differentiation, and function. TSC2 in the tuberous sclerosis complex potently regulates mTORC1 activation. CD8+ T-cells lacking TSC2 have constitutively enhanced mTORC1 activity and generate potent effector T cells; however sustained mTORC1 activation prevents generation of long-lived memory CD8+ T-cells. Here we show manipulating TSC2 at Ser1365 potently regulates activated but not basal mTORC1 signaling in T cells. Unlike non-stimulated TSC2 knockout cells, CD8+ T-cells expressing mutant TSC2-S1365A (SA) have normal basal mTORC1 activity. PKC and T-cell Receptor (TCR) stimulation induces TSC2 S1365 phosphorylation and preventing this with the SA mutation markedly increases mTORC1 activation and T-cell effector function. Consequently, CD8+ SA T-cells display greater effector responses while retaining their capacity to become long-lived memory T-cells. CD8+ SA T-cells also display enhanced effector function under hypoxic and acidic conditions. In murine and human solid-tumor models, CD8+ SA T-cells used as adoptive cell therapy have greater anti-tumor immunity than WT CD8+ T-cells. These findings reveal an upstream mechanism to regulate mTORC1 activity in T-cells. The TSC2-SA mutation enhances both T-cell effector function and long-term persistence/memory formation, supporting a novel approach to engineer better CAR-T cells to treat cancer.

## Introduction

The mammalian target of rapamycin (mTOR) is an evolutionarily conserved serine/threonine kinase which integrates cues from the environment to regulate cellular growth (1). In T-lymphocytes and throughout the immune system, mTOR is a potent regulator of cell activation, differentiation, and function (2, 3). This is achieved by multiple signaling pathways that prominently involve metabolic programming, cell growth, proliferation, and survival (4). MTOR signals via two multi-protein complexes, mTORC1 and mTORC2, that differentially regulate T cells (5). For CD4+ T cells, signaling downstream of mTORC1 promotes T_H_1 and T_H_17 differentiation while mTORC2 regulates T_H_2 differentiation (6, 7). Interestingly, antigen recognition in the absence of mTOR signaling leads to the generation of T regulatory cells (8, 9). For CD8+ T cells, mTORC1 activation is critical for effector cell generation and function (10, 11) while its inhibition promotes the generation of memory CD8+ T cells (12). mTORC2 ablation in CD8+ T cells also promotes memory T cell generation but does not inhibit effector T cells (10).

mTORC1 activity is tightly regulated upstream by the tuberous sclerosis complex (TSC) which consists of TSC1 (hamartin) and TSC2 (tuberin) (13, 14). TSC2 is a GTPase activating protein (GAP) for the small GTPase Ras homolog enriched in brain (Rheb) residing on the lysosomal surface along with the mTORC1 complex, and TSC is a constitutive inhibitor of mTORC1 in a Rheb-dependent manner. TSC2 activity is potently influenced by phosphorylation of multiple residues on TSC2 that are mediated by distinct kinases to alter mTORC1 activity based on the various environmental signals. Most prominently, the stimulation of the PI3K-PDK1-AKT-TSC2 axis by growth factors or T cell activation activates mTORC1 signaling through inhibitory phosphorylation of TSC2 at serine 939 to allow Rheb to activate mTORC1 through still unknown mechanisms (13). An equally potent TSC inhibitory effect is conferred by ERK1/2 phosphorylation at S540/S664 (15). Alternatively, increased AMP levels indicative of metabolic stress activates AMP activated protein kinase (AMPK) phosphorylating TSC2 at T1271/S1387 thus enhancing its ability to suppress mTORC1 signaling (16). Also, under environmental hypoxic stress, REDD1 (Regulated in development and DNA damage response-1) induction competitively binds to 14-3-3 proteins releasing the cytosolic sequestration of the TSC1/2 complex to inhibit mTORC1 activity (17).

Consistent with the role of mTORC1 in promoting effector CD8+ T cells, TSC2 KO (TSC2^-/-^) CD8+ T cells with constitutive elevated mTORC1 activity demonstrate enhanced effector function and display superior anti-tumor activity (10). However, TSC2^-/-^ CD8+ T cells fail to convert to long-lived memory T cells. Recently, phosphorylation of TSC2 at Serine 1365 (S1365 in mice; S1364 in human) was shown to be a critical site for the negative regulation of mTORC1 in stressed cardiac myocytes, fibroblasts, and intact hearts (18-20). Mutation of this site from serine to an alanine (S→A, SA) to prevent its phosphorylation, resulted in increased mTORC1 activation but only during states of pathological stress. Importantly, unlike TSC2^-/-^ cells which typically demonstrate constitutive mTORC1 activity, TSC2SA cells and mice with a SA-knock-in mutation did not display increased basal mTORC1 activity. Similar phenotypes were observed in heterozygous and homozygous SA-KI mice, indicating an autosomal dominant effect by possessing one mutant copy. Mutating the same serine to glutamic acid (TSC2SE) to create a phospho-mimetic (S→E, SE) also left basal mTORC1 activity unchanged, but greatly attenuated mTORC1 activation upon stress. MTORC1-stimulation pathways particularly regulated by the phospho-state of S1365 (or its genetic substitution with A or E) were mediated by mitogen activated kinases, including ERK1/2, whereas Akt activation of mTORC1 did not appear modified by S1365 status (19). While several kinases can result in phosphorylation of TSC2 S1365 including protein kinase C and cGMP-stimulated kinase-1 (cGK-1), we found the latter most relevant for suppressing mTORC1 co-activation in cardiomyocytes. The role of this signaling node in T-cells or other immune cells remains unknown.

Given these prior observations, we speculated phosphorylation of S1365 on TSC2 may also potently regulate activation-induced mTORC1 activity in T cells. Furthermore, we hypothesized that the SA silencing mutation at this site might promote enhanced effector CD8+ T cell generation due to increased mTORC1 activity immediately upon activation while retaining the ability to become long-lived memory T cells with mTORC1 activity returning to basal levels. Here, we reveal a novel role for S1365 in regulating mTORC1 activity in T cells that supports a novel approach to generate T cells with superior anti-tumor activity.

## Results

### TSC2 constitutively suppresses mTORC1 activity and inhibits T cell effector cell generation

In the canonical mTORC1 signaling pathway, environmental cues lead to PI3K and PDK1 activation which leads to the phosphorylation (T308) and activation of AKT which in turn leads to the phosphorylation and inactivation of TSC2 (S939). CD8+ T-cells activated by T-cell receptor (TCR, signal 1) engagement, PMA to PKC or interleukin-2 (IL-2), promote AKT and TSC2 phosphorylation and the downstream activation of mTORC1 as measured by phosphorylation of S6K1 and S6 (Figure 1A). Both PMA and TCR activation also lead to increased Akt and ERK1/2 phosphorylation. IL-2 but no IL-7 stimulation leads to strong phosphorylation of Akt but not ERK1/2, and this results in TSC2 phosphorylation at the Akt mediated site S939 to derepress mTORC1 inhibition by TSC suggesting different types of stimuli leads to differential signaling cascades. Genetic deletion of TSC2 in CD8+ T cells (T-TSC2^-/-^) increases constitutive baseline mTORC1 activity and also TCR-induced mTORC1 activity (Figure 1B) as we previously reported (10). Moreover, T-TSC2^-/-^ CD8+ T cells proliferate more readily in normal but also stressful acidic conditions emphasizing TSC2’s strong ability to sense environmental cues for modulating mTORC1 activity (Figure 1C). As we previously described, CD8+ T cells lacking TSC2 in response to an acute infection potently proliferate in vivo compared to WT T cells. To support our earlier study, TSC2 KO P14 CD8+ T cells recognizing LCMV were generated through CRISPR editing with naïve T cells (two different guides tested). We then co-adoptively transferred equal (1:1) numbers of Control or TSC2-KO edited P14 CD8+ T cells into naive WT recipient mice and tracked their response to acute LCMV Armstrong infection, measuring donor-cell frequency based on their surface congenic markers (Sup. Figure 1). On Day 8 of the peak acute response, TSC2 knockout CD8+ T cells again proliferated more robustly compared to control T cells (Fig. 1D). However, on Day 300 post infection, there were far fewer memory TSC2 KO CD8+ T versus control T cells. Thus, while the CD8+ T cell effector response is markedly enhanced through the genetic deletion of TSC2, sustained mTORC1 activity in the absence of antigen recognition inhibits the generation of long-term memory T cells. This highlights the key therapeutic limitation of genetically deleting TSC2 in CD8+ T cells due to the inability to repress mTORC1 signaling especially in the setting of adoptive cell therapy where memory potential is unquestionably required for better efficacy.

**Figure 1:**
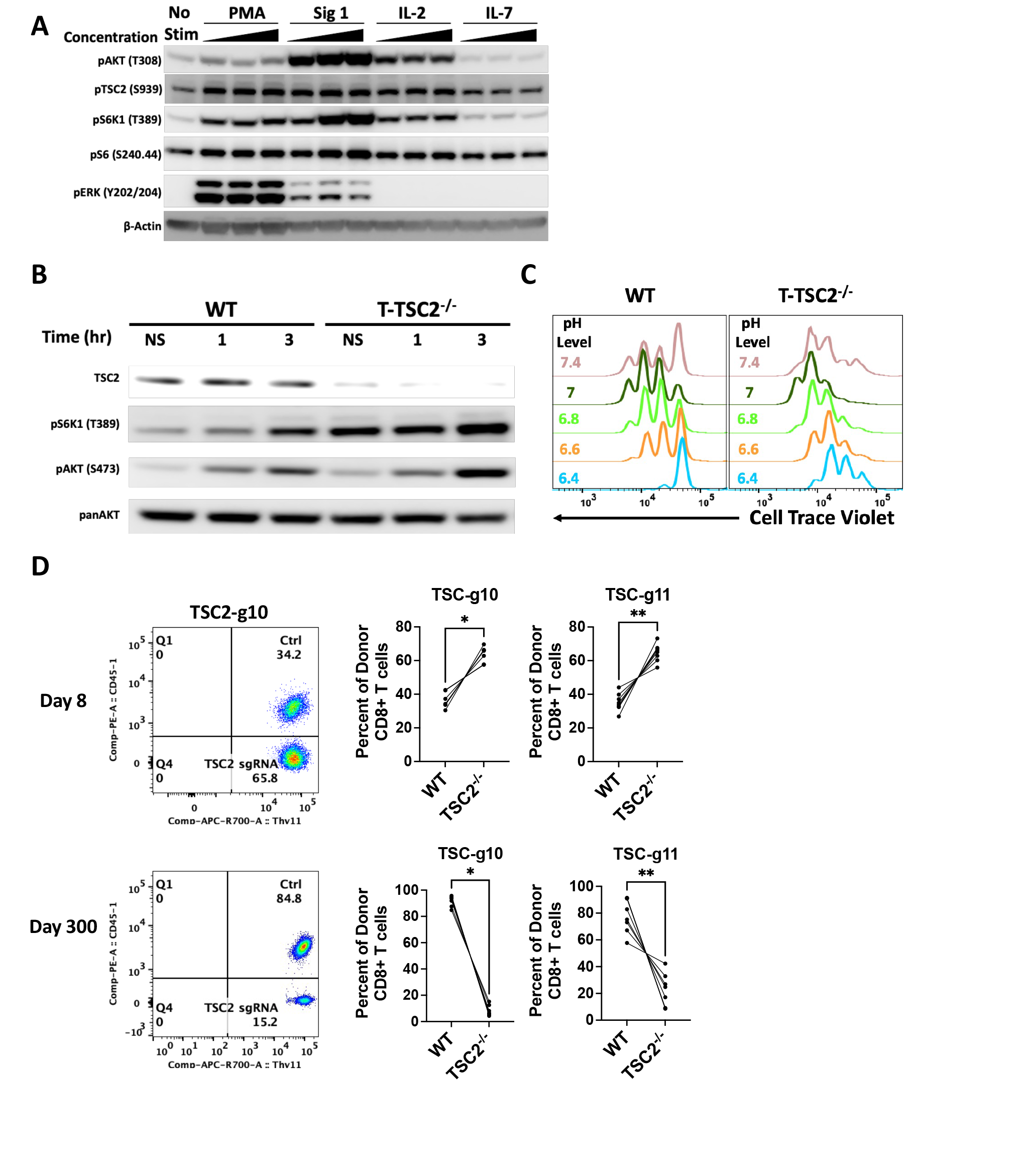
TSC2 is a central environmental hub that modulates mTORC1 activity and differentiation in T cells A) Immunoblot analysis of the PI3K-AKT-TSC2-mTORC1 pathway in resting T cells upon dose dependent stimulation of different stimuli for 30 minutes. B) WT and T-TSC2^−/−^ CD8+ T cells were stimulated with Signal 1 (TCR, αCD3) and Signal 2 (Co-Stim, αCD28) for 1 and 3 hours to assess mTORC1 and mTORC2 activity by immunoblot analysis; 0 hour indicates baseline no stim condition. C) Day 3 flow cytometry proliferation analysis of WT and T-TSC2^−/–^ CD8+ T cells stimulated and grown in different pH level media conditions. D) Different congenic naïve WT P14 (gp33+) CD8+ T cells were isolated and CRISPR edited with CAS9 and control sgRNA (Ctrl) or sgRNAs targeting TSC2 (g1 and g2). Ctrl and TSC2 sgRNA CD8+ T cells were mixed at 1:1 ratio and co-adoptively transferred into naïve WT recipients and infected with LCMV Armstrong for acute (Day 8) and memory analysis of donor CD8+ T cells via flow cytometry. *p<0.05, **p<0.01 paired t test. Data are representative of at least three independent experiments (A-C), two independent experiments (D).

### TSC2 S1365 is phosphorylated in T cells

Serine 1365 resides in a region of TSC2 containing multiple phosphorylation sites modified by other kinases that result in enhanced TSC suppression of mTORC1 (Figure 2A) (18). Inhibiting phosphorylation at this site by mutating it to an alanine (TSC2SA, termed SA mutant) led to a marked increase in stimulated mTORC1 activity in cardiac myocytes. Likewise, mutating this site to a glutamic acid, creating a phospho-mimetic (TSC2SE, termed SE mutant), led to a marked decrease in stimulated mTORC1 activity. Neither mutation affected basal mTORC1 activity but rather either enhanced (TSC2SA) or mitigated (TSC2SE) mTORC1 signaling in response to stimulation. Furthermore, in cardiac myocytes it was determined that cGK-1 was responsible for the selective phosphorylation of TSC2 at this site (18).

**Figure 2:**
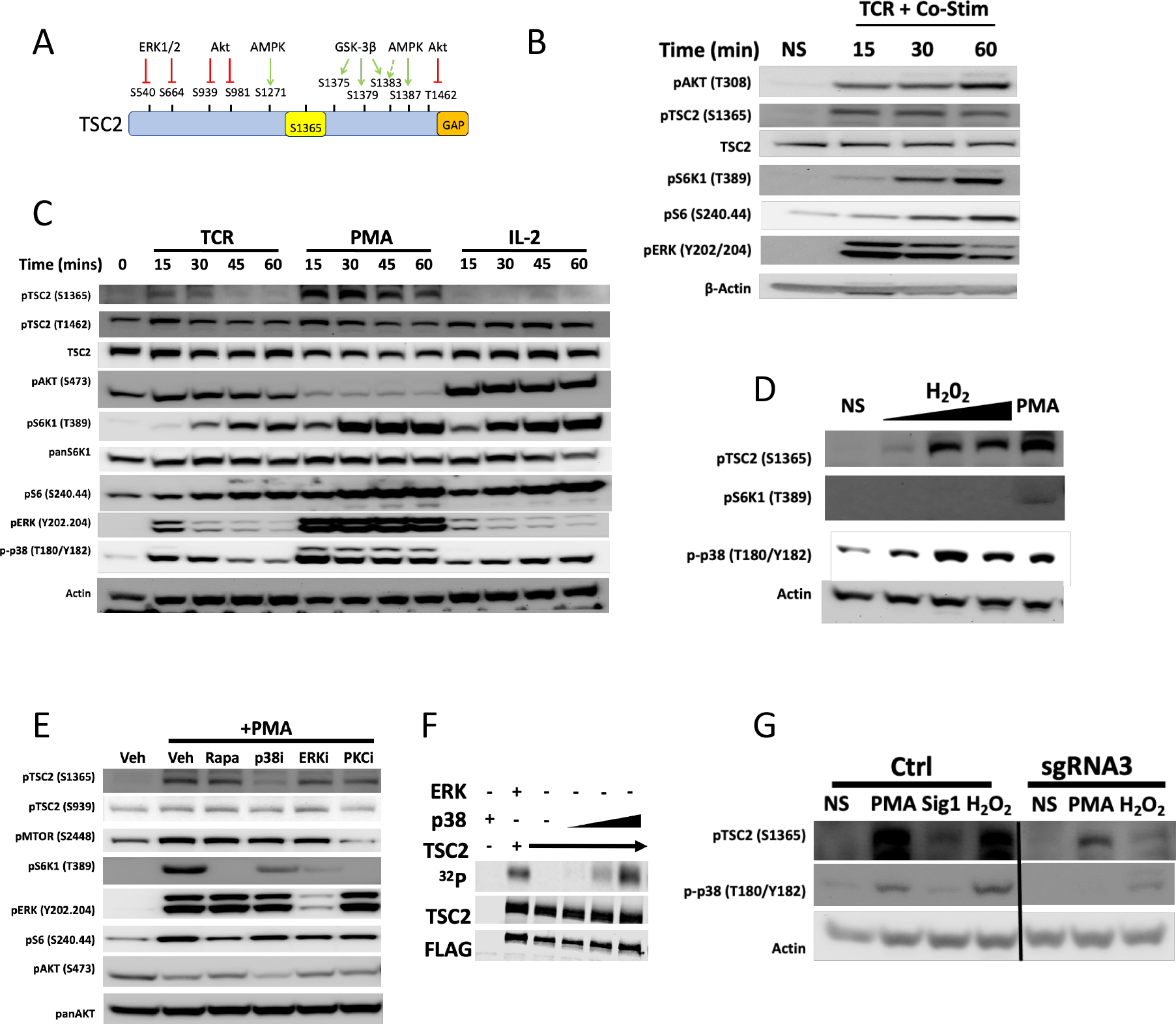
Identification of a novel TSC regulatory node (S1365) in T cells A) TSC2 protein map shows S1365 relative to other known phosphorylation sites for mouse and human protein. B) Immunoblot analysis of T cells stimulated with Signal 1 (TCR, αCD3) and Signal 2 (Co-Stim, αCD28) at indicated time points (minutes). TSC2 (S1365), PI3K (pAKT (T308)) and mTORC1 (pS6K1 (T389) and pS6 (S240.44)) activity were assessed. C) As in B, but stimulated with Signal 1 (TCR, αCD3) alone, PMA, or IL-2. D) T cells were exposed to increasing concentrations (.01mM, .1mM, 1mM) of H_2_O_2_ for 30 minutes and assayed by immunoblot analysis as indicated. PMA for 30 minutes is a positive control. E) T cells were stimulated with PMA in the presence of Vehicle (Veh), rapamycin, a p38 inhibitor (p38i), an ERK inhibitor (ERKi) and a PKC inhibitor (PKCi). The p38 inhibitor and to a lesser extent the PKC inhibitor blocked phosphorylation at TSC2(S1365). F) Flag tagged TSC2 protein was expressed in TSC2 KO HEK 293T cells and then immunoprecipitated with FLAG beads. Immune complexes were washed in lysis buffer and then in kinase buffer and incubated with ERK (positive control) or increasing amounts of purified active p38 and 2 μCi [γ− 32P]-ATP for 30 minutes. The reactions were stopped and prepped for autoradiography detection for 32P incorporation. G) Naïve T cells were transiently transfected with CAS9-sgRNA3 (p38-alpha) and subsequently activated with Signal 1 (TCR, αCD3) and Signal 2 (Co-Stim, αCD28) and expanded with IL-2 to generate activated T cells. On Day 7, viable CD8+ T cells were isolated and stimulated with PMA or TCR (Sig1) or exposed to .1mM H_2_O_2_ for 30 minutes. Data are representative of at least three independent experiments (B-E), two independent experiments (G).

Based on these findings in cardiac myocytes, we wondered whether TSC2 S1365 plays a biological role in regulating mTORC1 activity in T cells. First, we stimulated T cells with anti-CD3 (Signal 1) + anti-CD28 (Signal 2). Interestingly, as early as 15 minutes post T cell stimulation, we observed phosphorylation of TSC2 at S1365 (Figure 2B). Phosphorylation kinetics of TSC2 were consistent with TCR-induced AKT and S6K phosphorylation; that is, TCR-induced signals that promote the activation of mTORC1 signaling simultaneously promote the phosphorylation of a site on TSC2 that reigns in the response. This is analogous to increased pS1365 observed in cardiac myocytes and fibroblasts exposed to pathological growth factor stimuli and cardiac disease stress in vivo (18). As shown in Supplemental Figure 2A and 2B, pS1365 is not observed in T-cells expressing either the SA/SA, SE/SE mutation or TSC2 deletion, confirming specificity of the immunoblot antibody and signal despite many neighboring phosphorylation residues on TSC2 near S1365.

We next tested if other mTORC1 activators can also phosphorylate pS1365 in T cells. We found robust phosphorylation very early in response to PMA which directly activates PKC and subsequently downstream signaling cascades such ERK1/2 and p38, all pathways that are actively downstream of proximal TCR signaling (21) (Figure 2C). However, we did not observe this in response to IL-2 stimulation even at high concentrations, which canonically signals through the PI3K-AKT pathway (Sup. Figure 2C). That is, although like TCR and PMA activation, IL-2 leads to the activation of mTORC1 (and mTORC2 as measured by Akt S473 phosphorylation), IL-2 does not lead to phosphorylation of TSC2 S1365. IL-2 increased Akt phosphorylation of TSC2 (T1642) but did not activate ERK1/2, and as we recently found in other cell types, the latter but not former results in phosphorylation and mTORC-1 regulation by pS1365 (19). Corresponding S1364 phosphorylation was observed with both TCR-induced activation and PMA in human CD8+ T-cells (Sup. Figure 2D). We also found that exposure to hydrogen peroxide (H_2_O_2_) increased pS1365 in parallel with p38 activation (a known H_2_O_2_ target) (Figure 2D), though expectedly, H_2_O_2_ alone had no effect on mTORC1 activity (Figure 2D). Together, these data establish for the first time that in T cells, TSC2 S1365 is phosphorylated by both select extrinsic immune activating signals (TCR engagement but not IL-2) as well as extrinsic environmental cues such as H_2_O_2._

In cardiac myocytes, we first reported TSC2 S1365 is phosphorylated by cGK-1 (18). However, cGK-1 agonism with cGMP or inhibition (DT3) did not alter pS1365 in T cells (Sup. Figure 3A). Public databases for RNA expression (Immgen (22)) and proteomics (ImmPRes) showed cGK-1 mRNA and protein levels are not basally expressed in CD8+ T cells compared to other AGC kinases, nor in other immune cells other than mast cells (Sup Figure 3B and C). To assess alternative kinases that might be responsible for S1365 phosphorylation, T cells were stimulated with PMA and co-incubated with rapamycin to test if downstream mTORC1 factors contributed, or to a selective p38, ERK1/2, or PKC inhibitor. mTORC1 activity (pS6K1) was reduced by all inhibitors, but only p38 blockade also reduced pS1365 (Figure 2E). Using a radiolabeled ATP kinase assay with recombinant p38 and TSC2, we found a dependent increase in TSC2 phosphorylation by p38 itself demonstrating p38’s direct ability to regulate TSC2 (Figure 2F). Lastly, we used CRISPR to delete p38a in naïve T cells. Here, we also observed marked reduction of pS1365 in response to either PMA or H_2_O_2_ to activate p38 signaling (Figure 2G). Together, these data show that TSC2 S1365 is phosphorylated in T cells in response to T cell activation in a p38 dependent manner.

### TSC2 S1365 phospho-mutants regulate stimulated T cell function

To test the functional consequences of modifying TSC2 S1365 to a phospho-silenced (SA) or mimetic (SE) in T cells, we studied primary T-cells from global SA and SE KI mice as previously generated (18). First, we noted no significant differences in thymic T-cell populations comparing TSC2^WT^, TSC2^SA/SA^ or TSC2^SE/SE^ mice (Sup. Figure 4A). We also found peripheral CD4+ and CD8+ T cells percentages in the spleen were similar among these groups, whereas percentages of CD8+ T cells declined and CD4+ T cells increased in T-TSC2^-/-^ mice (Sup. Figure 4B-C). Importantly, consistent with our previously observed sustained mTORC1 activity even at rest (10), only T-TSC2^-/-^ CD8+ T-cells exhibited an elevated activation profile (CD62LloCD44hi) under basal conditions given their constitutive mTORC1 activity (Sup. Figure 4D).

We next isolated WT and SA/SA or SE/SE CD8+ T cells and stimulated them with anti-CD3 (TCR) + anti-CD28 (Co-Stim), measuring mTORC1 activity by pS6K1 (T389). Non-stimulated cells expressing SA/SA had no constitutive pS6K1 activity but showed amplified responses to TCR/Co-Stim (Figure 3A). By contrast, pS6K1 remained suppressed despite TCR/Co-Stim in T cells expressing the SE/SE mutation (Figure 3B). Importantly, basal mTORC1 activity measured by flow cytometry by pS6 (S240) was unaltered in either SA/SA or SE/SE, in marked contrast to an order of magnitude increase in T cells completely lacking TSC2 (Figure 3C). As we had previously found the SA mutation acts as an autosomal dominant, we also tested the impact of expressing a WT/SA mutation. Here too, we found marked augmentation of mTORC1 activation with TCR/Co-Stim (Figure 3D). This was further confirmed by flow cytometry showing resting WT, WT/SA, and SA/SA CD8+ T cells have minimal mTORC1 activity, but with TCR/Co-stim, both WT/SA and SA/SA T cells exhibit a 2-order of magnitude increase in pS6 phosphorylation (Figure 3E). Given the critical role for mTORC1 in CD8+ T cell proliferation, we next assessed proliferation 48 and 72hrs after T cell activation by monitoring cell trace dilution. We observed greater proliferation in SA/SA CD8+ T cells than WT/SA, and both more than WT T cells (Figure 3F). Lastly, we analyzed CD8+ T cell effector function based on of IFN-γ, TNFα, and IL-2 expression upon restimulation. Like the elevated proliferation and mTORC1 activity, WT/SA and SA/SA CD8+ T had increased effector function compared to WT T cells with the highest in SA/SA T cells (Figure 3G). In contrast, SE/SE CD8+T cells had reduced effector function compared to both WT and SA/SA CD8+ T cells (Sup. Figure 4E). Together, these data reveal the importance of the phosphorylation of this single amino acid, TSC2 S1365 to regulate T cell activation-induced mTORC1 activity and consequently proliferation and effector function.

**Figure 3:**
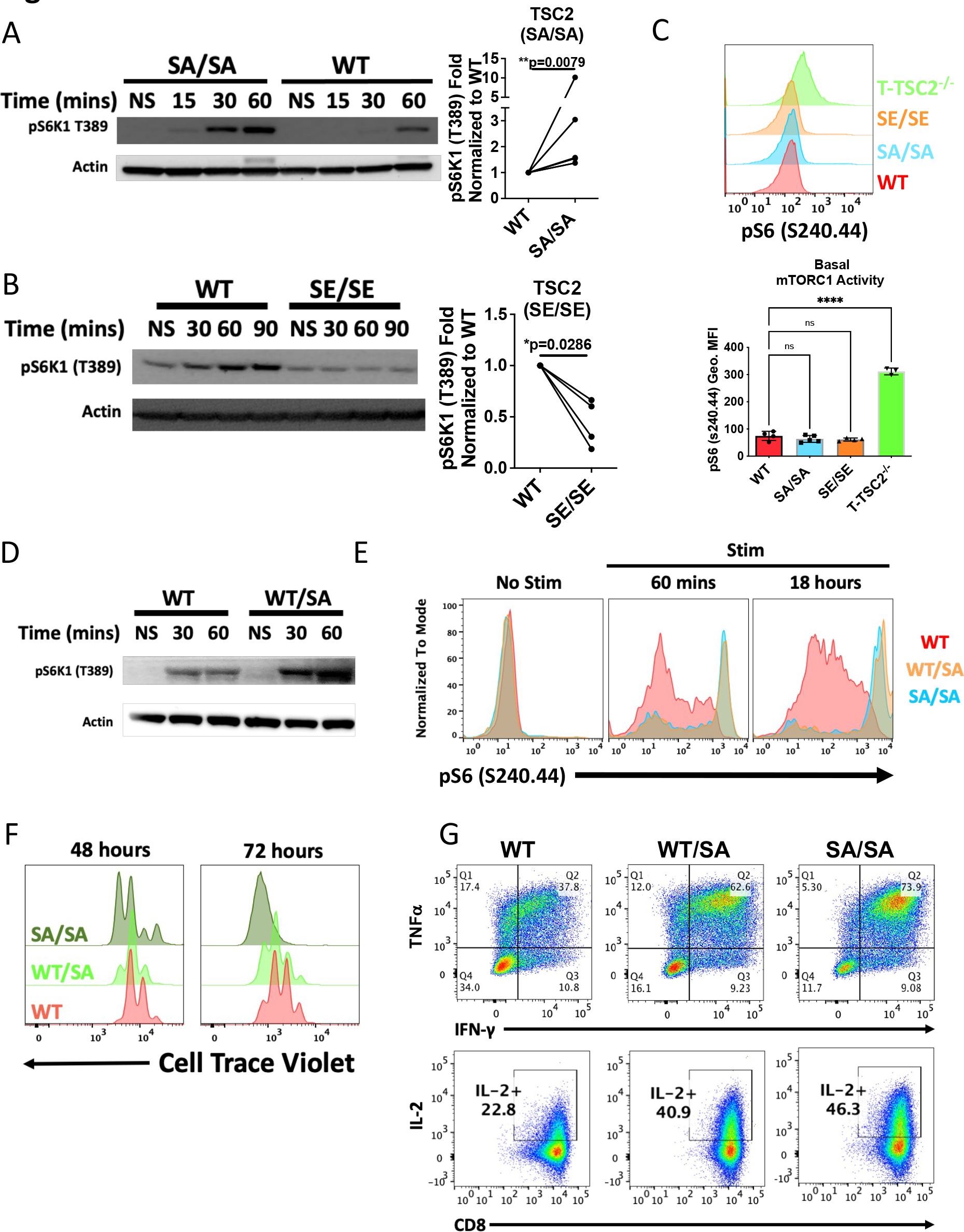
Mutating TSC2 at S1365 augments mTORC1 activity and function in murine CD8+ T cells upon stimulation A) Immunoblot analysis of mTORC1 activity in resting WT and mutant TSC2SA/SA (S→A, serine to alanine) CD8+ T cells stimulated with Signal 1 (TCR, αCD3) and Signal 2 (Co-Stim, αCD28) for indicated time points. B) As in A but with WT and TSCSE (S→E, serine to glutamic acid) CD8+ T cells T cells. C) mTORC1 activity was assessed by intracellular staining for mTORC1 activity via phospho-S6 levels in naïve WT, TSC2SA, TSC2SE, and T-TSC2^-/-^ CD8+ T cells Note, the unstimulated levels (NS) of mTORC1 activity for the mutant CD8+T cells are equivalent to that of the WT CD8+ T cells unlike T-TSC2^-/-^ CD8+ T cells. D) As in A, but using TSC2WT/SA CD8+ T cells. E) Naïve WT, WT/SA, and SA/SA T cells were stimulated for 60 minutes and 18 hours and mTORC1 activity was assessed by intracellular staining for mTORC1 activity via phospho-S6 levels. F) WT, WT/SA and SA/SA mutant CD8+ T cells were stimulated to analyze cell proliferation (CTV) with flow cytometry on Day 2 and Day 3 post-activation. G) As in E, CD8+ T cells were expanded in IL-2 to generate effector cytotoxic lymphocytes for cytokine analysis of IFN-γ, TNFa, and IL-2 upon re-stimulation. Statistical graphs on right depict normalized fold change of mTORC1 activity based on densitometry between TSC2 mutant to WT CD8+ T cells at last time point in independent experiments. *p<0.05, **p<0.01, ****p<0.0001. one-way ANOVA with Tukey’s multiple comparisons test (C). Data are representative of at least 3 independent experiments.

### TSC2(SA) T cells demonstrate enhanced effector and memory generation in vivo

Given the strong in vitro observations, we next tested the effector response of TSC2(SA) CD8+ T cells in vivo. Differentially Thy marked, WT or SA/SA-OTI (TCR recognizing OVA) CD8+ T cells were co-adoptively transferred 1:1 into naïve WT recipients, that were then infected with Listeria-OVA to induce an acute immune response (Sup. Figure 5A). On Day 8, we analyzed the relative percent of WT vs SA/SA expressing CD8+ T cells in the spleen. As we observed in vitro, we found a much higher percent of SA/SA CD8+ T cells to WT T cells demonstrating a strong ability to proliferate in response to infection (Fig 4A). Interestingly, we did not observe any phenotypic differences (terminal (KLRG1+) versus memory precursors (CD127+)) between WT and SA/SA CD8+ T cells though we observed enhanced effector function in SA/SA CD8+ T cells (Figure 4B-C). SA/SA CD8+ T cells are alike to TSC2^-/-^ CD8+ T cells in that both display greater proliferation and effector function, but TSC2^-/-^ CD8+ T cells are also prone to be terminally differentiated, a critical contrasting phenotype between the two genotypes. We then compared WT/SA OTI CD8+ T cells to both WT and SA/SA T cells in a triple co-adoptive setting into the same WT host. Again, we found more homozygous and heterozygous TSC2(SA) mutant CD8+ T cells versus WT CD8+ T cells (Figure 4D). Thus, even a single copy of the TSC2(SA) mutation promoted enhanced T cell proliferative responses in vivo. We also found greater numbers of SA/SA CD8+ T cells at later time points (6 weeks) post-infection indicating a strong ability to persist long-term as memory T cells in strong contrast to TSC2^-/-^ CD8+ T cells (Figure 1D and 4E).

**Figure 4:**
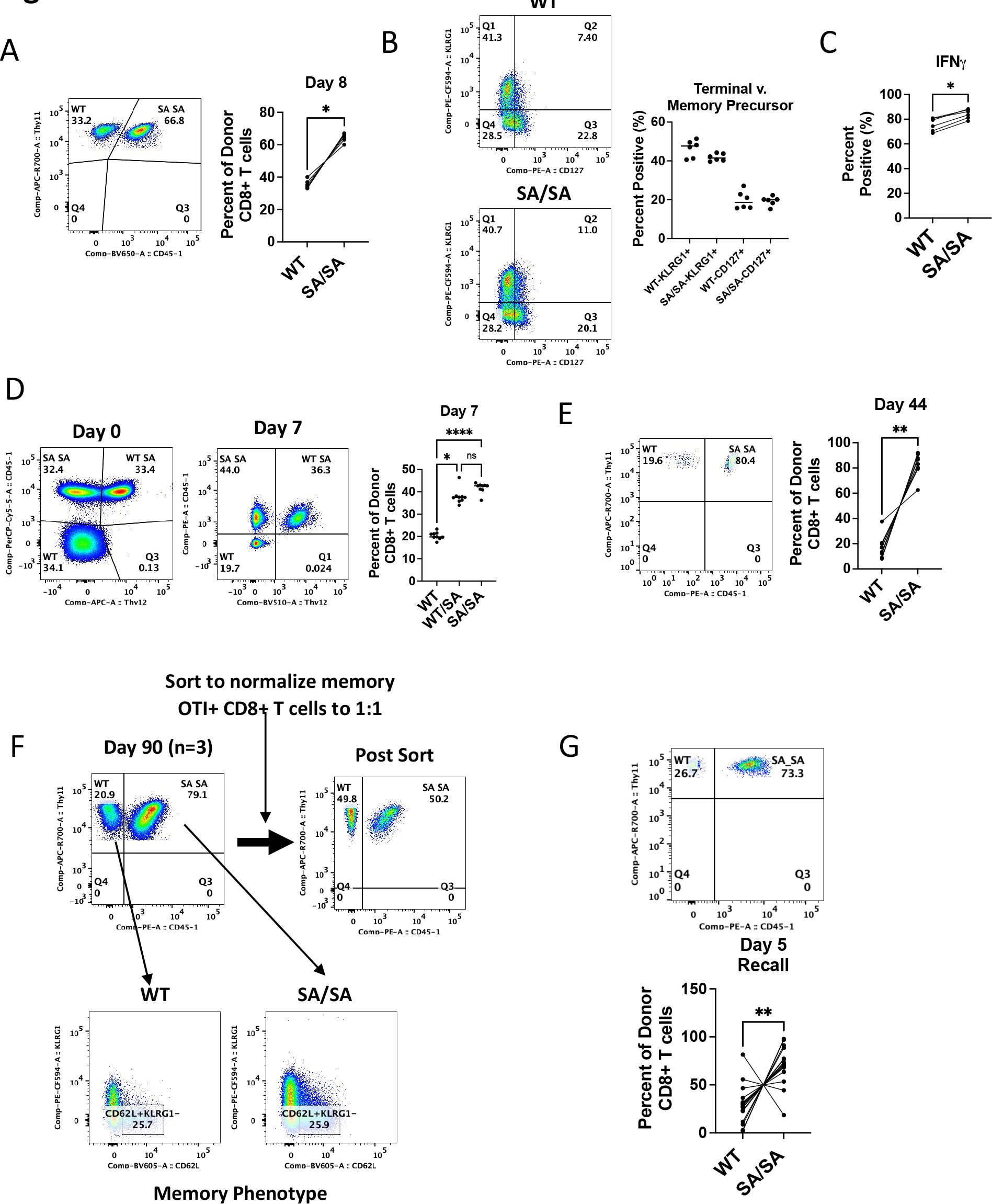
Mutating TSC2 at S1365 (SA) promotes strong CD8+ T cell effector response while preserving memory formation and recall capacity WT and mutant TSC2 (SA) transgenic CD8+ T cells were co-adoptively transferred into naïve WT hosts followed with acute pathogen infection. A) Flow cytometry plot of transferred CD8+ T cells (left) and summary data (right) showing percent of WT vs SA/SA genotype from donor population 8 days after exposure to Listeria-OVA. B-C) Phenotypic and functional analysis of donor WT and mutant TSC2SA CD8+ T cells during acute infection. D) Triple co-adoptive transfer of WT, mutant heterozygous (WT/SA) and homozygous (SA/SA) TSC2 CD8+ T cells with listeria-OVA infection to look at numbers on Day 7 of acute response by flow cytometry. E) Percent present of donor WT and mutant TSC2 SA CD8+ T cells after 44 days (memory phase). F) Percent of memory WT and mutant TSC2SA OTI CD8+ T cells in spleen (left, n=3). Equal number of memory donor CD8+ T cells were sorted and again co-adoptively transferred into naïve WT recipients to assess memory recall ability on 1:1 basis upon infection. Bottom: memory characterization of memory CD8+ T cells. G) From F, percent of donor CD8+T cells after Day 5 upon recall. *p<0.05, **p<0.01, ****p<0.0001. A paired T-test was performed for statistical analysis. Data are representative of at least 3 independent experiments.

To test if we generated more bona fide memory CD8+ T cells with the SA mutation unlike TSC2^-/-^ CD8+ T-cells (10), we repeated the co-adoptive transfer study using WT and SA/SA memory CD8+ T cells injected into new WT recipients to assess their ability to functionally recall, a cardinal tenet of memory T cells. We again found more memory SA/SA than WT CD8+ T cells 90 days post infection (Figure 4F). These resting memory CD8+ T cells were then sorted to co-transfer (1:1) again into naïve WT hosts that were then re-infected. Importantly, the WT and SA/SA memory CD8+ T-cells demonstrated no significant differences in their central memory phenotype based on CD62L+KLRG1-expression indicating both genotypes were similar going in prior to rechallenge (Figure 4F, bottom). However, on Day 5 post re-challenge, we again found more SA/SA compared to WT CD8+ T cells, demonstrating their enhanced recall ability (Figure 4G). Similar results were obtained in studies using WT/SA CD8+ T-cells and using LCMV Armstrong as another infection model (Sup. Figure 5B-D). Thus, the SA mutation at S1365 supports CD8+ T cells to markedly enhance T cell effector generation without compromising memory formation and recall capacity in response to infection in vivo, which significantly differs from the TSC2^-/-^ CD8+ T cell phenotype.

### TSC2(SA) T cells display enhanced activation under conditions of cellular stress

The tumor microenvironment is generally considered both hypoxic and acidic thus hampering mTORC1 activity and anti-tumor immune responses (23). Since TSC2 is a central metabolic and stress hub in mammalian cells, we hypothesized that the SA mutation in CD8+ T cells might provide an intrinsic advantage to support better proliferation and effector function under environmental stress. Activation of mTORC1 reflected by pS6K1 induced by either PMA or TCR stimulation was substantially decreased under hypoxia (1-2% O_2_) compared to normoxic conditions (Figure 5A). Next, we stimulated WT and WT/SA CD8+ T-cells under these conditions and monitored proliferation. Even with one copy of this mutation in CD8+ T cells, we observed a strong ability for WT/SA CD8+ T cells to proliferate under hypoxia (Figure 5B) and generate more effector cytokines upon rechallenge (Fig. 5C) in both normoxic and hypoxic conditions. MTORC1 activation is acutely reduced by acidosis as shown Figure 5D. This is accompanied by activation of p38 and correspondingly more TSC2 S1365 phosphorylation which suggests its augmentation of TSC2 activity helps suppress mTORC1 in low pH conditions. We also observed strong TSC2 S1365 phosphorylation in human T cells in response H_2_O_2_ and low pH (Sup. Figure 6). Since TSC2^-/-^ CD8+ T cells can tolerate low pH conditions (Figure 1C), we tested whether blocking S1365 phosphorylation with the SA mutant would augment mTORC1 activity in acidic conditions. As shown in Figure 5E, in lower pH conditions, greater mTORC1 activity was seen in CD8+ T cells expressing either the WT/SA and SA/SA mutant copy (Figure 5E). Furthermore, we observed increased IFN-γ production in SA/SA CD8+ T cells when compared to WT CD8+ T cells over a broad range of pH (Figure 5F). These results demonstrate that S1365 is a potent signaling node that responds in stressful environmental conditions, and that its inhibition by the SA mutation enhances CD8+ T cell mTORC1 activation and effector function despite hypoxia or acidosis.

**Figure 5:**
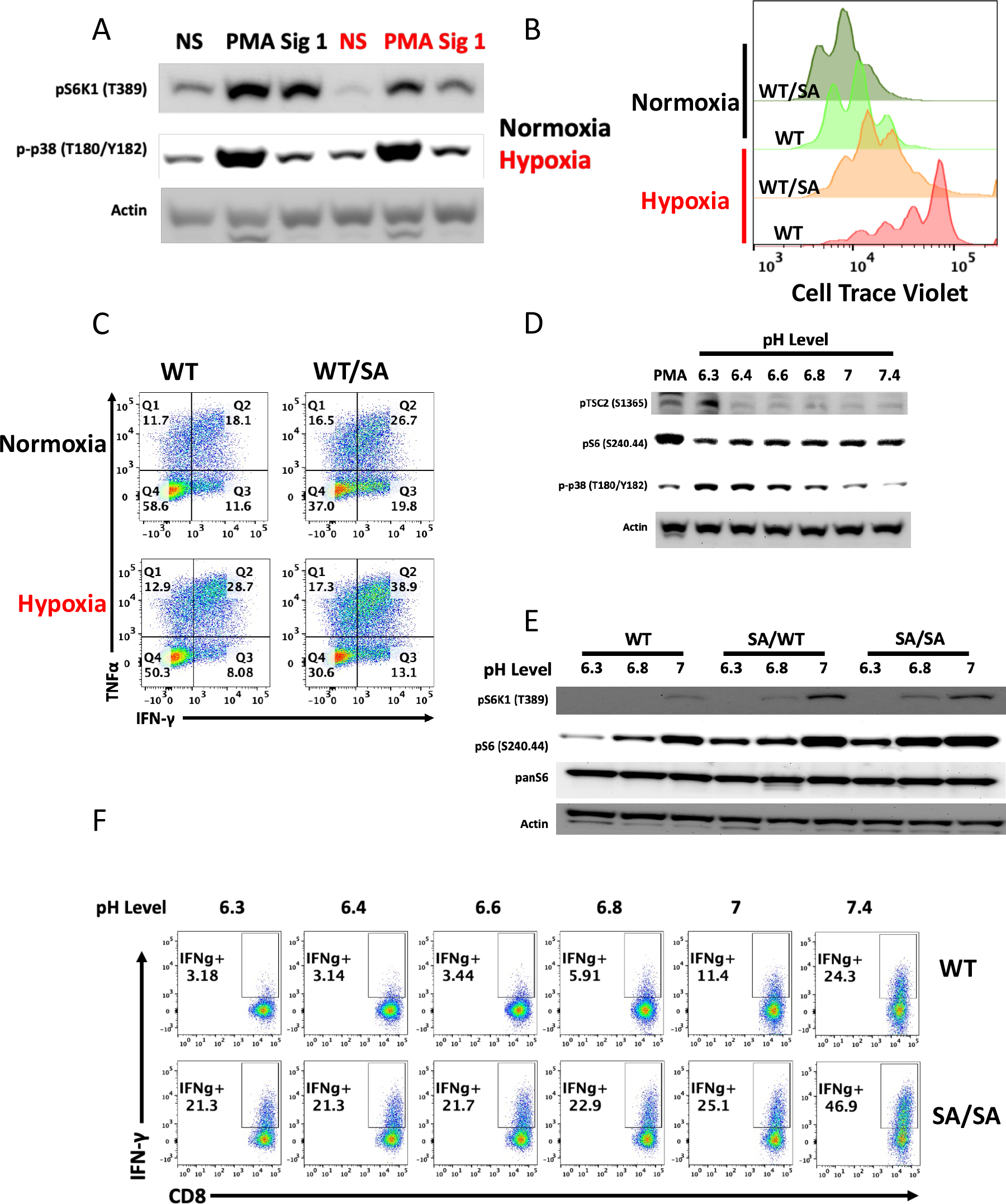
SA mutation in CD8+ T cells amplifies mTORC1 activation under cellular stress. A) Resting CD8+ T cells were stimulated with either PMA or TCR (Signal 1) in either normoxic or hypoxic (2% O_2_) conditions for 15 minutes and assayed by immunoblot analysis as indicated. B) Mutant TSC2SA CD8+ T cells are more resistant to hypoxia: Cell trace violet (CTV) labelled naïve WT and mutant TSC2 (WT/SA) OTI CD8+ T cells were stimulated with peptide for three days in normoxic or hypoxic (2% O_2_) to assess proliferation by CTV dilution via flow cytometry. C) IL-2 generated cytotoxic WT and mutant TSC2SA CD8+ T cells re-challenged overnight for cytokine analysis in normoxic or hypoxic (2% O_2_) by flow cytometry. D) Activated WT, mutant TSC2 WT/SA and SA/SA CD8+ T cells were exposed to various pH level media conditions for 90 minutes and assayed by immunoblot analysis for pTSC2 and mTORC1 activity. E) Effector WT or TSC2 SA/SA CD8+ T cells were stimulated with PMA and Ionomycin in various pH level media to assess IFN-γ via flow cytometry. Data are representative of at least three independent experiments except (F) with two.

### TSC2(SA) mutation enhances anti-tumor adoptive cellular therapy

Having established that TSC2 S1365 regulates CD8+ T cell effector and memory generation as well as CD8+ T cell responses under hypoxic and acidic conditions, we next tested if CD8+ T cells harboring the TSC2(SA) mutation would demonstrate superior anti-tumor activity. To this end, we first chose the B16-OVA model which is relatively resistant to adoptive cellular therapy. Transfer of activated WT OTI CD8+ T cells had limited impact on tumor growth or long-term survival, whereas transfer of either WT/SA or SA/SA OTI CD8+ T cells significantly inhibited tumor growth and enhanced survival (Figure 6A). Based on earlier in vitro experiments, we predicted CD8+ T cells with the SA mutation were better able to tolerate the harsher tumor microenvironment. To test this, we co-adoptively transferred equivalent numbers of activated WT and SA OTI CD8+ T cells into B16-OVA tumor bearing mice. Four days after T cell transfer, tumors were processed and analyzed to assess the frequency of WT and SA mutant donor CD8+ T cells in the tumor (TILs). Within the donor CD8+ T cells, SA mutant T cells comprised the vast majority (∼90%) compared to WT T cells (Figure 6B). WT/SA CD8+ T-cells were phenotypically less exhausted based on lower surface expression levels of the exhaustion markers PD1 and LAG3 versus WT CD8+ T cells with enhanced cytokine IFN-γ expression (Figure 6C).

We next tested if SA expression enhanced the efficacy of adoptively transferred CD8+ T cells using a murine CD19 CAR-T cell model treated with WT or SA/SA CD8+ T cells (Sup. Figure 7A) (24). As in the OVA-OTI model, SA/SA CD19 CAR-T cells also significantly suppressed tumor growth (Figure 6D). We again performed a co-adoptive transfer experiment using equal number of WT and WT/SA CD19 CAR-CD8+ T cells to assess the TILs, and once more found WT/SA T cells were significantly increased in the tumor yet not the draining lymph node (Fig 6E, Sup. Figure 7B-C). Consistent with our previous data (Figure 6B), SA mutant TSC2 CAR CD8+ T cells are better able to survive and to proliferate in the solid tumor microenvironment.

**Figure 6:**
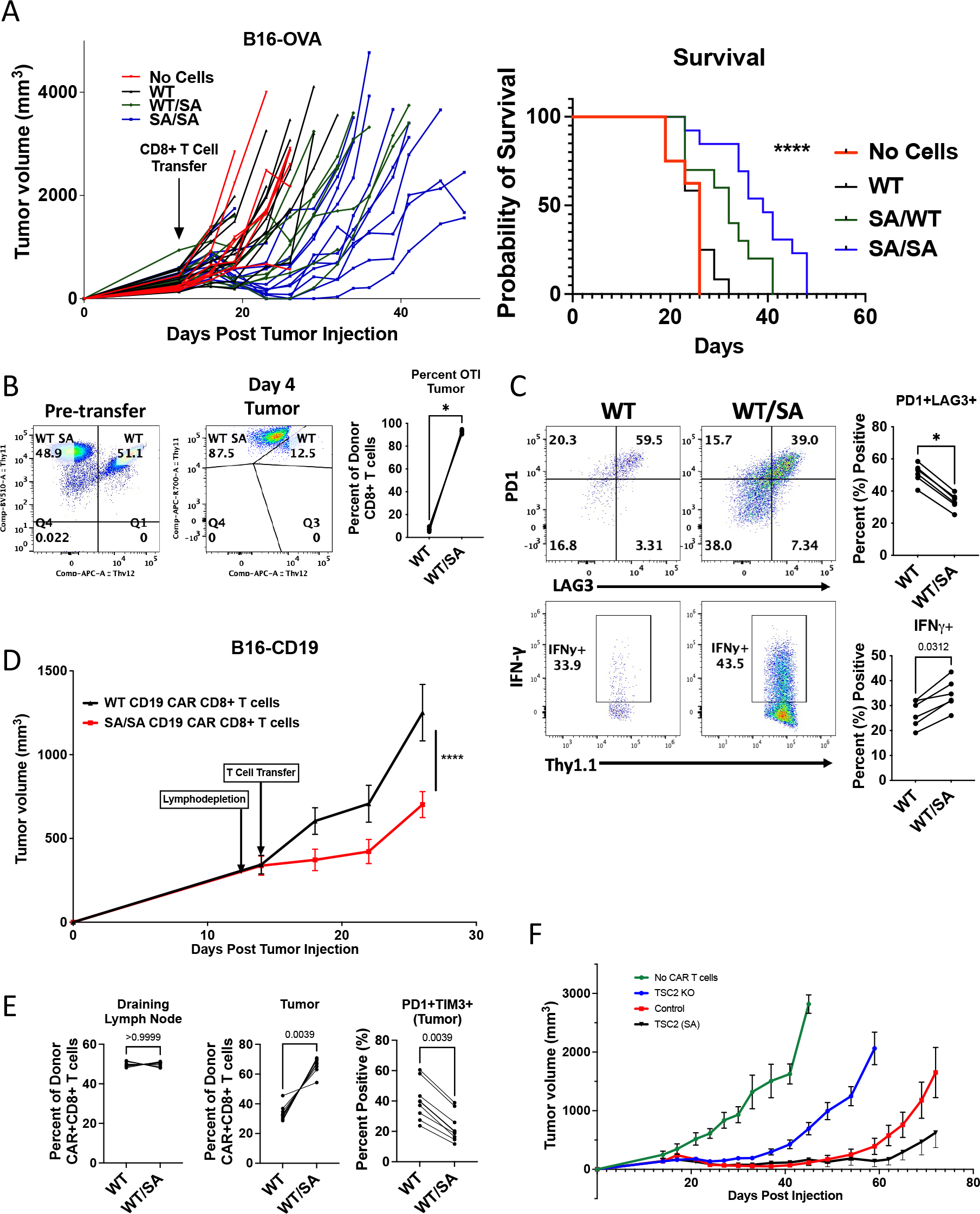
TSC2 SA mutant CD8+ T cells promote strong anti-tumor immunity with adoptive T cell therapy. A) In vitro activated WT, mutant TSC2 SA OTI CD8+ T cells were injected into WT recipients that had received B16-OVA cells 11 days prior. Tumor volume was assessed every 2 to 3 days. Survival was assessed. Mice that received mutant TSC2 SA CD8+ T cells had enhanced survival compared with all other treatments, as determined by multiple comparisons Mantel-Cox tests (right). B) Equal number of WT and TSC2 (SA) heterozygous OTI CD8 co-transferred into the same tumor bearing host (left). Flow cytometry analysis of tumor infiltration CD8+ T cells between donor WT and mutant TSC2SA CD8+ T cells (right). C) Flow cytometry analysis of exhaustion profile (top) and function (bottom) of donor WT and mutant TSC2SA CD8+ T cells. Numbers in quadrants indicate percent cells in each (throughout). D) As in 6A but using a murine CD19 CAR T cell model towards B16 tumors expressing CD19. Growth curves until first sacrifice. E) As in 6B, but analyzing donor CAR T cells in draining lymph node (dLN) and tumor on Day 4 and Day 8. F) As in 6A but using human CD70 CAR T cells towards CD70 expressing human tumor in NSG mice. A paired T-test was performed for statistical analysis (*p<0.05, **p<0.01, ***p<0.001, ****p<0.0001). Data are representative of at least 3-6 independent experiments (A-C), 3 independent experiments (D-E), and 2 independent experiments (F)

Lastly, to further test the clinical potential of engineering this mutation in human T cells, we successfully generated human TSC^-/-^ CD70 CAR-T cells and SA mutant CD70 CAR-T cells (Sup Figure 8). We then compared their anti-tumor efficacy in a CAR T cell model to CD70 expressing human non-small cell lung cancer (NSCLC) tumors in NSG mice. We again found SA CAR T cells resulted in better tumor control for a longer duration versus WT CAR T cells and TSC2^-/-^ CD70 CAR-T cells as the key comparison group (Fig 6F, Sup Figure 9A). In vitro, we observed increased IFN-γ production in SA CAR T cells when compared to Control T cells under different low to high pH conditions but no changes in cytotoxic lytic capacity (Sup. Figure 9B, C). Interestingly, while all group of CAR-T cells were able to initially suppress tumor growth, the TSC^-/-^ were the least effective in controlling tumor growth likely given their poor persistence and memory potential. This is consistent with the loss of TSC^-/-^ T cells over time and provides a striking example of how mutating a single node (SA) on TSC2 substantially differs from deleting TSC2.

## Discussion

MTOR has emerged as an important integrator of signals dictating T cell differentiation and function. A critical component of its ability to regulate immune cells is via metabolism (9), as mTORC1 activation promotes T cell effector function by upregulating glycolytic metabolism (10, 25). Likewise, inhibition of mTORC1 enhances long-lived memory CD8+ T cell generation by inhibiting glycolytic programs and promoting oxidative phosphorylation. In addition, the role of mTOR in regulating cytokine signaling and the expression of canonical transcription factors have been shown to contribute the ability of signaling downstream of mTOR to regulate T cell differentiation and function (26). It is this complex coordination of T-cell functionality that has made manipulation of mTORC1 therapeutically challenging. mTORC1 inhibitors such as rapamycin or suppressors of upstream activation by PI3K and Akt have long been a target for cancer, but they also suppress T-cell immune effector function. Here we have shown that a single mutation in TSC2 at position S1365 (SA) to block its intrinsic phosphorylation increases mTORC1 activation upon T cell stimulation leading to enhanced effector function. Importantly, mutation at this site does not result in constitutive mTORC1 activation but selectively enhances activity upon stimulation. For T cells, this translates to the ability of the SA mutation to promote the generation of effector T cells resulting in a concomitant increase in long-lived memory T cell generation. This results in adoptive T-cell therapy exhibiting both enhanced tumor-suppression and persistence with less T-cell exhaustion. This constellation of features has been keenly sought after to improve CAR-T and other adoptive cell therapies, and the present findings support use of gene-editing to generate a TSC2 SA mutation as part of the strategy.

While mTORC1 is regulated by a multiplicity of signals, including glucose, lipids, amino acids, and signaling kinases, it is the latter that prominently converge on TSC2 to modify mTORC1 complex signaling (13, 14, 27-29). Akt, ERK1/2, p38, RSK1 kinase activation and 14-3-3 binding all relieve TSC2 inhibition of mTORC1, whereas AMP-activated kinase and GSK3β enhance TSC2-mediated mTORC1 suppression. Despite this central role, modification of TSC2 has been previously difficult to leverage as a means of selectively manipulating mTORC1 activity. First, most kinase-regulated changes engage multiple residues to generate the effect, making gene editing more complex. Second, mutagenesis of these sites to prevent or mimic phosphorylation has led to altered constitutive mTORC1 activity. Serine 1365 on TSC2 is unique in this regard, as we find minimal basal impact with either modification, yet very potent directionally opposite changes upon mTORC1 stimulation. This was first revealed in fibroblasts and cardiac myocytes (18) and is now recapitulated in T cells. It is this feature that differentiates the SA mutant from TSC2^-/-^ T-cells, the latter showing similar acute amplified effector function upon stimulation but poor long-term persistence and memory. However, the SA mutation preserves overall TSC2 functionality but amplifies effector function and proliferation, however, it only does so with selective activation, so persistence and memory formation are also improved.

In contrast to cardiac myocytes, in T cells, phosphorylation of TSC2 S1365 is not mediated by cGK-1 but rather by p38 MAPK. PKC may also achieve this though still via a downstream p38 mechanism. Importantly p38 and PKC are activated upon TCR stimulation along with ERK1/2 and Akt. That S1365 is intrinsically phosphorylated in conjunction with PKC stimulation or TCR engagement supports this as an intrinsic negative regulator to modify these mTORC1 activation pathways. While not explored in detail here, our recent study found S1365 phosphorylated along with MAPK but not Akt activation, and this mirrored whether phospho-mutants altered mTORC1 stimulation (19). We also found disparate phosphorylation between TCR and IL-2 stimuli, and while not shown here, suspect S1365 modifications will have analogous selectivity to T cell input signaling and thus cytokine control. Importantly, TCR induced mTORC1 activity is regulated by S1365 status affording a means to improve CAR T cell activation in response to low tumor antigen expression.

Tumor-infiltrating lymphocytes in different tumor models showed substantial bias to T-cells harboring the SA mutation over WT. This could reflect metabolic changes and/or synthetic changes coupled to mTORC1 activation that enabled SA T-cells to better survive and proliferate within the tumor microenvironment. Our in vitro data showing both enhanced proliferation and cytokine generation of SA T cells despite hypoxia or acidosis is also consistent with this. Expression of the SA mutation in the KI mouse has also been shown to enhance cardiac protection against ischemic damage by a mechanism linked to greater glucose versus fatty acid metabolism (20). SA-T cells also consistently exhibited less markers of exhaustion, supporting an underlying capacity to regulate mTORC1 activation dependent on the level of co-stimulation.

The current study findings have potential clinical implications. Immunotherapy in the form of Adoptive Cellular Therapy (ACT) has emerged as a potent means to treat cancer (30). ACT provided by CAR-T cells is an FDA approved treatment for several hematologic malignancies. However, there remain many challenges including the lack of long-lived memory of the CAR-T cells, and less success in the treatment of solid tumors (31). This may be due in part to the hostile tumor microenvironment, and to that end, creating CAR-T cells with a TSC2(SA) mutation may help circumvent this limitation by improving effector function and long-term survival even in hypoxic and/or acidic microenvironments. Furthermore, use of these modified T-cells may promote better persistence of CAR-T cells that can be quickly re-activated and expanded to prevent relapse. This also suggests potential synergy with check-point inhibitors, as these therapies can re-awaken quiescent T-cells but not those that are terminally exhausted. Finally, while the technology to create such genetically engineered CAR-T is now available, the fact that the TSC2(SA) mutation acts as an autosomal dominant, with generally similar impact from a hetero- and homozygote mutation, suggests that mutation efficiency of even 50% should be quite effective. As engineering of CAR-T cells becomes more diverse and facile, targeted base editing or even expression of a mutant protein possibly linked to a controllable inducer are but two such examples of exploiting our findings clinically. These and further translational studies to test the impact of TSC2(SA) should define their value for immuno-oncology therapies.

## Materials and Methods

### Mice

Six- to ten-week-old male or female mice were used for performing all the experiments in this study. NSG, C57BL/6J, CD4 Cre, CD8+ OTI (OVA TCR), CD90.1/Thy1.1+, CD45.1 mice were obtained from Jackson Laboratories and CD4+ 5CC7 (PCC TCR) from Taconic and bred in-house. P14 (gp33 TCR) were kindly provided by David A. Hildeman, University of Cincinnati. Mice with knock-in mutations at S1365 (Serine to Alanine (SA) and Serine to Glutamic Acid (SE)) were as first generated on a C57BL/6J background (18) and used in the study. Mice with loxP flanked TSC2 alleles were generated by the laboratory of Michael Gambello (University of Texas Health Science Center at Houston, Houston, Texas) and bred to CD4 Cre. Genotyping determined by respective protocols. No empirical test was performed for choosing sample size prior to experiments. No randomization of samples or animals was used nor were investigators blinded throughout the study.

### Antibodies and Reagents

Antibodies against the following proteins were purchased from BD Biosciences: CD8a (53-6.7), CD90.1 (OX-7), CD90.2 (53-2.1), CD45.1 (A20). Antibodies against PD1 (29F.1A12), LAG3 (C9B7W), CD127 (A7R34), CD62L (MEL-14), were purchased from Biolegend. Fixable viability dye ef480 was purchased from eBioscience. The following antibodies were purchased from Cell Signaling: TSC2 (D93F12), pTSC2 (S939), pS6 (S240/244, D68F8), AKT (11E7), pAKT (S473, D9E), pAKT (T308, D25E6), p4E-BP1 (T37/46, 236B4), pMTOR (S2448, D9C2), mTOR (7C10), pERK (T202/Y204, D13.14.4E), pFoxo1 (T24/32, 9464). p-p38 MAPK (Thr180/Tyr182) (D3F9), β-Actin (13E5). pTSC2 (S1365) (120718, NovoPro Labs). Class-I peptide (SIINFEKL, OVA I) was purchased from AnaSpec. Fc Block (2.4G2) and stimulatory in vivo plus grade anti-CD3 (2C11) and anti-CD28 (37.51) were purchased from BioXcell. Cell Proliferation Dye-eFluor450 was purchased from eBiosciences. Rapamycin was purchased from LC laboratories. All small molecule inhibitors were purchased from Cayman Chemicals and used at the following concentrations: ERK (U-0126, 10uM), PKC (Calphostin C, 1uM), p38 (SB 203580, 5uM), mTORC1 (Rapamycin, 1uM). Secondary fluorophore conjugated antibodies were purchased from Invitrogen: anti-rabbit Alexa Fluor647. IL-2 (10ng/mL) and IL-7 were purchased from Peprotech. PMA (50ng/mL), Ionomycin (100ng/mL), H_2_O_2_ was purchased from Sigma. Golgi Stop was purchased from BD bioscience.

### Radiolabeled ^33^P TSC2 Kinase Assay

TSC2 KO HEK cells were transduced with adFLAG-TSC2. Samples were collected and underwent immunoprecipitation for FLAG. The kinase assay was performed while TSC2 was in the immunocomplex on the magnetic beads, with either TSC2 alone (negative control), activated ERK2 kinase + TSC2 (positive control), p38 alone (negative control), and increasing p38 dose combined with TSC2. Samples were then eluted from the beads by boiling in sample loading buffer and subjected to SDS-PAGE and Western blotting for TSC2 and FLAG and imaging by film for ^33^P incorporation.

### Activating T cells in vitro

Naïve resting splenocytes from spleens and lymph nodes were combined for all experiments. In summary, single cell suspensions were created by mashing organs through 70uM filter. Splenocytes from WT, TSC2 KO, and TSC2 SA mice were stimulated with soluble anti-CD3 (3ug/mL) or isolated CD8+ T cells (Negative Selection, Biolgend) with plate-bound anti-CD3 (5ug/ml) and soluble anti-CD28 (2ug/mL) for 48 hrs then expanded in IL-2 (10ng/mL) for 4-5 days to generate resting previously activated CD8+ T cells. WT or TSC2 mutant (SA or SE) OTI CD8+ T cells (final 5e6/mL) were stimulated with OVA peptide (100ng/mL) for 48 hrs then expanded and fed with fresh media and IL-2 (10ng/mL) daily for 4-5 days to generate resting previously activated effector CD8+ T cells for further functional or signaling assays. 5CC7 CD4+ PCC Transgenic T cells were stimulated with 5ug/mL PCC peptide for 48hrs then expanded in IL-2 (10ng/mL) for 4-5 days to generate previously activated CD4+ T cells for signaling analysis. Cell proliferation of OTI CD8+ T cells with peptide (100ng/mL) stimulation was monitored with cell trace violet (CTV, eBio) between 48-72 hrs.

### T cell activation or stress induced signaling

Naïve T cells from spleens and lymph nodes were combined for all experiments. In summary, single cell suspensions were created by mashing organs through 70uM filter and CD8+ T cells were isolated using negative selection isolation. Naïve or resting IL-2 expanded T cells from WT, TSC2 knockout (T-TSC2^-/-^), mutant TSC2 SA, and mutant TSC2 SE mice were stimulated with cross-linked with Armenian Hamster IgG, anti-CD3 (3ug/mL) anti-CD28 (2ug/mL), PMA, H_2_0_2_, or other activation stimulus at indicated time points for flash freezing to assess signaling via immunoblotting. Primary T cell cultures were maintained in RPMI-1640 media (Corning 10-040-CV) with 10% FBS (Gemini Bioproducts), 2 mM L-glutamine (Corning), 10 mM HEPES (Corning 25-060-CI), gentamycin (50ug/mL, Quality Biological), non-essential amino acids (100x, Gibco), and beta-mercaptoethanol (50uM, Sigma) in standard humidified 5% CO2, 37° C tissue culture incubators. Experimental culture conditions described below.

### Assessing T cell mTOR activity via flow cytometry

Primary splenocytes were derived and stimulated as detailed above. Following stimulation, splenocytes were fixed with 2% PFA for 10 minutes at 37 deg then washed two times with PBS. Cells were permeabilized with ice cold 90% methanol for 20 minutes at -20 deg. Cells were washed three times with 1% FBS/PBS (staining solution). Next, cells were stained with anti-CD4 (1:500) and anti-CD8 (1:500) and anti p-S6 (S240.44) (1:2000) in staining solution for 45 minutes at room temperature. Cells were then washed two times with staining solution before stained with Rabbit IgG AF647 (1:500) for 30 minutes at room temperature. Cells were then washed two times afterwards. Gates were set appropriately with the aid of unstimulated and secondary alone controls. All experiments were performed on a BD FACS Calibur, LSR II or Aria II and analyzed using FlowJo software analysis.

### Intracellular Cytokine Stimulation

Viable T cells were enriched with ficoll gradient then stimulated with PMA/Ionomycin (4-5 hours) or platebound anti-CD3 (1ug/mL) and soluble anti-CD28 (2ug/mL) (14-16 hours) with GolgiStop to assess cytokine production. Cells were collected then stained with viability dye and surface staining for 20 minutes at 4 deg. Next, cells were washed and fixed with 100uL of BD CytoFix/Perm kit for 30minutes at room temperature, washed with BD 1x PermWash buffer, and stained with intracellular cytokine staining in 1x BD PermWash for 45 minutes at room temperature before analyzed on flow cytometer.

### Low pH Experiments

For experimental manipulation of pH, CD8+T Cells from WT, TSC2 KO, TSC2 SA were stimulated in RPMI-1640 (Sigma R1383 with 11.1 mM glucose restored) supplemented with 10% FBS, 2 mM L-glutamine, gentamycin (50ug/mL), and beta-mercaptoethanol (50uM) in which the bicarbonate-CO2 buffering was replaced with 25 mM PIPES and 25 mM HEPES in atmospheric CO2 as above. Cultures were maintained at 37° C in a humidified incubator. When prepared, a slightly concentrated media was split into multiple volumes before adjusting pH to target values, bringing to volume, and sterilizing by filtering, ensuring identical media composition in all regards other than pH. pH of stored media was frequently monitored to guard against slow drift and ensure correct record of experimental conditions (32).

### LCMV, Vaccinia-OVA, and Listeria (LM)-OVA CD8+ T Cell Adoptive Transfer Model

For assessing acute and memory responses in vivo, different congenically marked isolated CD8+ OTI or P14 T cells were mixed at equal ratios (1E3 cells of each population) and co-adoptively transferred into wild-type hosts. Recipient mice were shortly later infected with double attenuated Listeria monocytogenes (5×106 cfu/mouse, i.v.), Vaccinia expressing ovalbumin antigen (VacOVA) (1×106 pfu/mouse, i.v.) LCMV Armstrong (2×10^5^ pfu/mouse, i.p.) LCMV Amstrong was kindly provided by Susan Kaech. Spleens were collected 6-8 days post-infection. Blood and spleens were collected at indicated time points. When indicated, resting memory mice were rechallenged using Vaccinia virus expressing OVA (VacOVA, 1×106 pfu/mouse, i.v.). For the secondary adoptive transfer experiment, resting memory splenocytes were harvested after naive adoptive transfer and infection. Cells were sorted for equal donor CD8+ T cells, and again co-adoptively transferred into naïve hosts for secondary recall with pathogen.

### B16-OVA and B16-CD19 Adoptive T Cell Therapy Model

For the B16 adoptive T cell therapy models, naive C57BL/6J WT mice received a s.q. injection of 2.5 × 10^5^ B16-OVA melanoma cells (gift of Hyam Levitsky) cultured in vitro under OVA selection media containing 400 μg/ml G418 (Life technologies) or 5 × 10^5^ B16-OVA-CD19 (kindly provided by Anjana Rao(24)). B16-OVA Model: 11 days after tumor inoculation, mice received an adoptive transfer of 7.5 × 10^5^ activated WT or mutant TSC2 SA OTI CD8+ T cells derived from splenocytes, which had been stimulated in vitro with SIINFEKL peptide (100ng/mL) for 48 hours and expanded in IL-2 (10ng/mL) for an additional 48hr hours. On Day 4 post-activation, cells were subjected to ficoll gradient to enrich for viable CD8+ T cells. TSC2 WT or SA mutant CD8+ OTI T cells were transferred into tumor bearing mice for tumor outgrowth experiments. Mice were randomized into groups on day of therapy. B16-OVA-CD19 Model: mice were lymphodepleted with 200ug Cyclophosphamide (IP, Sigma) one day before receiving CD19 CAR+ CD8+ T cells. Tumor volume was calculated using the formula for the prolate ellipsoid, (L × W2)/2, where L represents length and is the longer of the 2 measurements and W represents width. Tumor burden was assessed every 2 to 4 days by measuring length and width of tumor. Mice were sacrificed when tumors exceeded 2×2cm, necrotic, or mice experienced visible signs of discomfort. To analyze tumor infiltrating T cells (TILs), a 1:1 ratio of cells was mixed to analyze infiltrate of CD8+ T cells on Day 4-8 post-transfer.

### Retroviral transduction of CD19 CAR CD8+ T cells

In brief, polyclonal isolated WT or mutant TSC2 SA CD8+ T cells were stimulated for 24 hours with plate-bound anti-CD3 (5ug/mL) and soluble anti-CD28 (2ug/mL) with human IL-2 (100units/mL). CD19 CAR+ retroviral transductions were performed in six-well non-tissue treated coated plates with 20ug/mL retronectin (Takeda). Fresh virus of (MSCV-myc-CAR-2A-Thy1.1, Anjana Rao, Addgene: 127890) containing chimeric antigen receptor (CAR) was made using Platinum-E (Plat-E) Retroviral Packaging Cell Line (Cell Biolabs). Fresh virus with human IL-2 was spin fected onto retronectin coated plates according to protocol (33156338). After spinfection of virus, activated T cells were slowly layered on top of the virus and briefly spun for two additional days on virus coated plates for viral transduction. On Day 3, T cells were collected and expanded in fresh media with 100units hIL-2/mL for 4 additional days before transferring into B16-OVA-CD19 tumor bearing hosts. CD19 CAR efficiency was assessed by Thy1.1 surface staining. Equal CAR+ CD8+ T cells were transferred into tumor bearing mice by normalizing with flow cytometry for single transfer (efficacy) and co-adoptive experiments (TILs).

### Tumor Infiltrating Lymphocyte (TIL) Harvest

Four days after CD8+ OTI or CD19 CAR-T cell transfer, tumors were harvested from mice and digested in 2 mg/ml collagenase I (Life Technologies) with DNase I (Roche) in RPMI 1640 supplemented with 2% FBS. Tumors were digested with constant rotation at 37 deg for 30 minutes followed by quenching with EDTA. Cells were then filtered and processed into single cell suspensions. Cells were then stained with Abs for subsequent flow analysis of transferred CD8+ T cells defined by congenic markers from tumor bearing host.

### Immunoblot Analysis

For immunoblot analysis, T cells were harvested by centrifugation and resuspended in ice-cold lysis buffer (20 mM Tris (pH 7.5), 150 mM NaCl, 1 mM EDTA, 1 mM EGTA, 1% Triton X-100, 2.5 mM sodium pyrophosphate, 1 mM β-glycerolphosphate (glycerol-2-phosphate), 1 mM sodium orthovanadate, 1 mM PMSF, 1× protease inhibitors (Roche, Basel, Switzerland)) and lysed at 4 °C for 30 min. Lysates were cleared of debris by high speed centrifugation. Equal protein mass from each condition was mixed with 4× LDS buffer (Invitrogen, Carlsbad, CA) and boiled for 10 min. Lysates were then loaded into NuPAGE gels (4-12% Bis–Tris gels, Invitrogen) and run at 150 V for 90 min. Protein was transferred to polyvinylidene fluoride (PVDF) membranes with transfer buffer (1× NuPAGE Transfer Buffer (Invitrogen) with 20% methanol) at 30 V for 90 min. Membranes were blocked in 5% nonfat dry milk (NFDM) for 60 min, washed briefly with Tris-buffered saline + 0.1% Tween-20 (TBST) and probed with primary antibody in 4% NFDM in TBST overnight at 4°C. Membranes were washed with TBST three times for 10 min and probed with secondary antibody conjugated to HRP in NFDM. Membranes were washed 2 times in TBST for 5 min, and then once in Tris-buffered saline once for 5 min. Enhanced chemiluminescent picoplus substrate (ThermoFisher) was used to detect HRP-labeled antibodies. Blots were developed using a Biospectrum Multispectrum Imaging System and images were acquired and analyzed using VisionWorks, LS Image Acquisition and Analysis software (UVP, Upland, CA).

Human CD70 CAR T cells: Western Blot assay was performed with normalized protein concentrations of CAR-T cell pellet lysates, (Lysis Buffer R0278, Sigma-Aldrich) in the presence of Halt Protease and Phosphatase Inhibitor (Thermo Scientific #78442). Equal total protein concentration is assayed, with WESTM equipment, with detection of TSC2 (Cell Signalling Technology, Tuberin/TSC2, catalogue # D93F12, XP® Rabbit mAb) and GAPDH (14C10, Cell Signaling Technology catalogue # 2118). TSC2 expression was normalized to GAPDH via signal peak area.

### CRISPR/Cas9 RNP system of naïve T cells

Modified single guide RNAs (sgRNAs) were designed and synthesized by Synthego. Ribonucleoproteins (RNPs) were prepared by incubating sgRNAs and Cas9 nuclease (Integrated DNA Technologies, IDT) for 10 min at room temperature. For the delivery of RNPs, isolated WT P14 CD8+ T cells were washed with PBS and mixed with RNPs by using P3 Primary Cell 4D-NucleofectorTM X Kit (Lonza) immediately prior to electroporation (Lonza 4D-nucleofactorTM core unit, program DN100). Electroporated T cells were recovered and washed with T cell culture media. Cells were activated with plate-bound anti-CD3 and soluble anti-CD28 and expanded in IL-2 for subsequent immunoblot analysis. The sequences of sgRNAs are as followed:

Ctrl: 5’-GCACUACCAGAGCUAACUCA-3’;

g1: 5’- CCCUUGUGUUUCCAGGUACC-3’;

g2: 5’- GAGAUGGGUCACCAGGUACC-3’;

g3: 5’- GUACCUGGUGACCCAUCUCA-3’.

Human CAR-T TSC2-SA/SA Generation and Protocol

Guide RNA was designed to disrupt the TSC-2 gene near S1363 using homologous recombination to change serine to an alanine. Point mutations were also introduced to the donor template to stop the Cas9/guide complex from cutting the donor template and mutated gene. Homology arms were designed to the sequences flanking the mutated sequence to direct the homology directed repair.

WT sequence GTTGGCAGG GGCATCCCCATCGAGCGAGTCGTCTCCTCGGAGGGT GGCCGG

donor template GTTGGCAGG GGCATtCCaATtGAaCGgGTtGTggctTCtGAaGGT GGCCGG WT sequence GIPIERVVSSEG

donor template GIPIERVVASEG

Guide used

Spacer TSC-4 CATCGAGCGAGTCGTCTCCTCGG

Guide sequence mC*mA*mU*CGAGCGAGUCGUCUCCUguuuuagagcuagaaauagcaaguuaaaauaaggcuaguccguu aucaacuugaaaaaguggcaccgagucggugcmU*mU*mU*U

mX = 2’O-methyl (M)

X* = 2’-ribo 3’-phosphorothioate (S)

Where X = nucleotide

### CAR T-cell production

CAR T cell constructs were synthesized and cloned into an AAV6 plasmid backbone. All CAR construct included a CD8 transmembrane domain in tandem with an intracellular 4-1BB costimulatory and CD3ζ signaling domain. Gene-editing and cell preparation was performed using standard techniques as described in detail elsewhere [29]. Briefly, human peripheral blood mononuclear cells (PBMCs) were thawed, and the T cells were activated with conjugated CD3/CD28 agonists for 3 days in T-cell media containing human serum, IL-2 and IL-7. After activation, the T cells were electroporated with Cas9 protein and sgRNAs targeting the TRAC and B2M loci or TRAC, B2M, and CD70 with or without TSC2 loci and subsequently transduced with a recombinant AAV6 vector containing donor template DNA for insertion of the CD70 CAR construct, with or without TSC2 S1364A Δ. Following electroporation and transduction, the CAR T cells were expanded for 7 days in T-cell media containing human serum, IL-2, and IL-7. These cells were frozen in Biolife solution CryoStor CS10 Freeze Media (Fisher Scientific NC9930384) and transferred to storage in liquid nitrogen prior to use in assays.

### T-cell assays

T-cell assays for activity, proliferation and cytotoxicity have been described in detail elsewhere [30]. Briefly, in coculture experiments, T cells were incubated with Daudi target cells at an effector to target ratio (E:T) of 0.5:1, 1:1 and 2:1 for 20 hours. Cell-free supernatants from cells were subsequently analyzed for cytokine expression using a Luminex array (Luminex Corp, FLEXMAP 3D) according to manufacturers’ instructions. Expression of surface markers were either taken at baseline or after a period of coculture, and then subjected to flow cytometric analysis. Antigens were specifically stained using the following antibody clones for flow cytometry where indicated: CCR7 (CD197, Biolegend); CD45RA (H1100, Biolegend). For proliferation, cells were counted every 1 to 3 days. Percent specific lysis for cell lysis determination may be calculated from live cell number, target cells pre-stained with specific marker (efluor670 (Invitrogen™ eBioscience™ Cell Proliferation Dye eFluor™ 670, catalogue # 50-246-095), normalized with flow cytometry counting beads (CountBright beads (Absolute Counting Beads for flow cytometry, Life technologies catalogue # C36950).

### TIDE Analysis of Genome Editing

TIDE analysis was performed as follows; isolate DNA of CAR-T cell pellets with DNEasy Blood & Tissue Kit (Qiagen catalogue # 69506), perform PCR amplification with appropriate forward and reverse primers to TSC2 S1364A Δ insertion and then submit PRC products for sequencing (Genewiz, Cambridge, MA). On receipt of sequencing data, sequences are analyzed relative to the TSC2 S1364A Δ sequence, with Tsunami software, to determine the % identical or aberrant sequences, insertions and deletions present in the samples. Additional SnapGene sequence alignment was performed with Blast 2 BlastSearch to confirm alignment.

### Statistical Analysis

All graphs and statistical analysis were performed using GraphPad Prism software (v.7 and 8). A p value less than 0.05 was considered statistically significant. Error bars represent mean ± standard deviation.

### Study Approval

Mice were maintained and studied in accordance with protocols approval by the Johns Hopkins University Institutional Animal Care and Use Committee.

## AUTHOR CONTRIBUTIONS

C.H.P, J.S., D.A.K and J.D.P designed and oversaw the study; C.H.P., Y.D., B.D.E, and M.J.R, L.M.B, D.B.H, performed experiments and data analysis. J.W. helped with mouse genotyping and colony maintenance. D.A.K and J.D.P acquired funding. C.H.P, D.A.K and J.D.P wrote the manuscript.

## ACKNOWLEDGEMENTS

We kindly thank Powell lab members for helpful discussion, Anjana Rao for human CD19 expressing B16 cells and murine CD19 CAR construct, and Ada Tam and Lee Blosser for cell sorting assistance. This work benefitted from data assembled by the ImmGen consortium and The Immunological Proteomic Resource (ImmPRes). This work was supported by the National Institutes of Health (T32HL007227 to C.H.P.; R01AI156274, R01AI07761, P41EB028239-01 to JDP, R35HL135827 to DAK, AHA fellowship award and 18CDA34110140 to MJR, F31-HL143905 to BLD-E) and The Bloomberg∼Kimmel Institute for Cancer Immunotherapy.

## DECLARATION OF INTERESTS

J.D.P. is a cofounder and equity holder of Dracen Pharmaceuticals. L.M.B, D.B.H, and J.S. are current employees of CRISPR Therapeutics. C.H.P. and J.D.P. are current employees of Calico LLC. Authors CHP, BLD-E, MJR, DAK, and JDP have filed patents for the use of TSC2 mutations at Serine 1364 and Serine 1365 (human) to treat disease.

## Notes

### Summary of Updates

Figure 3 - minor change to make the flow cytometry images clearer in lower panels, and Figure 4 - arrows identifying source of cell analysis in lower panels of Panel 4F.

