## Supplemental figures and legends for "Signaling Node at TSC2 S1365 Potently Regulates T-Cell Differentiation and Improves Adoptive Cellular Cancer Therapy"

**Sup. Figure 1**

A

Day 0-Pre-transfer check

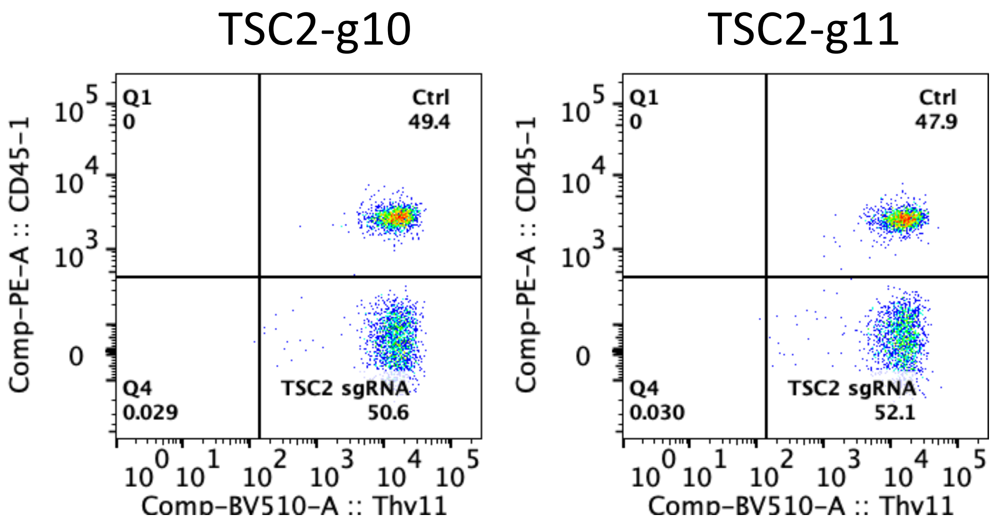


B

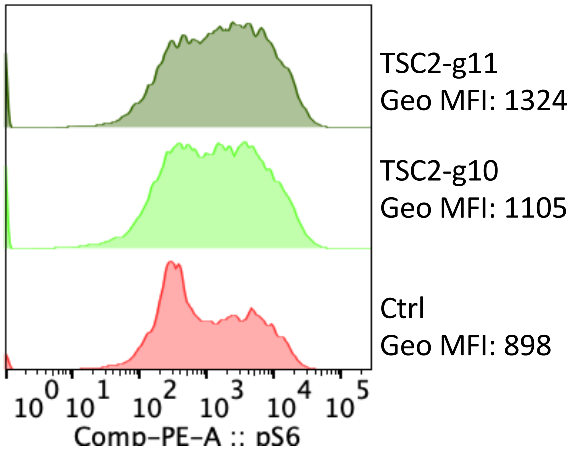

Supplemental Figure 1: TSC2 is a central environmental hub that modulates mTORC1 activity and differentiation in T cells. A) As in Figure 1, pre-transfer check of co-adoptive Ctrl and TSC2 sgRNA edited donor P14 CD8+ T cells. B) mTORC1 activity was assessed by intracellular staining for mTORC1 activity via phospho-S6 levels in Ctrl sgRNA and TSC2s sgRNA CD8+ T cells from Figure 1 to show enhanced mTORC1 activity upon TSC2 deletion. Representative of two independent experiments.

**Sup. Figure 2**

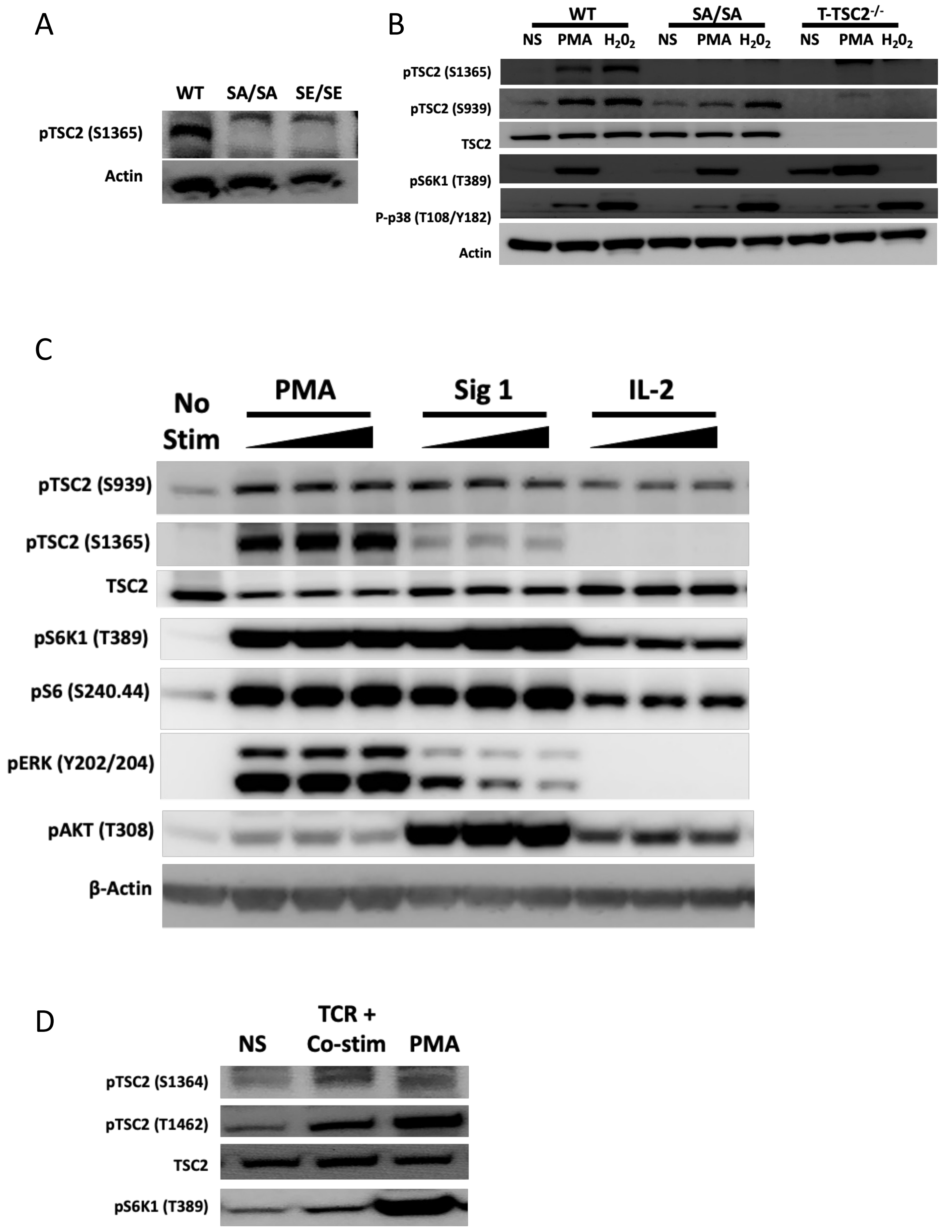
A
WT
SA/SA
SE/SE
pTSC2 (S1365)
Actin
B
WT
SA/SA
T-TSC2$^{-/-}$
NS
PMA
$H_20_2$
NS
PMA
$H_20_2$
NS
PMA
$H_20_2$
pTSC2 (S1365)
pTSC2 (S939)
TSC2
pS6K1 (T389)
P-p38 (T108/Y182)
Actin
C
No Stim
PMA
Sig 1
IL-2
pTSC2 (S939)
pTSC2 (S1365)
TSC2
pS6K1 (T389)
pS6 (S240.44)
pERK (Y202/204)
pAKT (T308)
β-Actin
D
NS
TCR + Co-stim
PMA
pTSC2 (S1364)
pTSC2 (T1462)
TSC2
pS6K1 (T389)

Supplemental Figure 2: pTSC2 (S1365) is active in T cells upon activation. A) Antibody validation using immunoblot analysis using PMA stimulated (30mins) WT, TSC2 SA/SA and SE/SE T cells. B) Additional antibody validation using immunoblot analysis using PMA or $H_2O_2$ stimulation (30mins) in WT, TSC2 SA/SA and TSC2$^{-/-}$ T cells. C) Immunoblot analysis of T cells stimulated with Signal 1 (TCR, αCD3) alone, PMA, or IL-2 in dose increasing manner for 30 minutes. TSC2 (S1365), PI3K (pAKT (T308)) and mTORC1 (pS6K1 (T389) and pS6 (S240.44)) activity were assessed. D) Previously activated human T cells were re-stimulated with Signal 1 (TCR, αCD3) and Signal 2 (Co-Stim, αCD28) or PMA for 30 minutes for immunoblot analysis to assess pTSC2 and mTORC1 activity. Data are representative of one experiment (A-B), two independent experiments (C-D).

### Sup. Figure 3

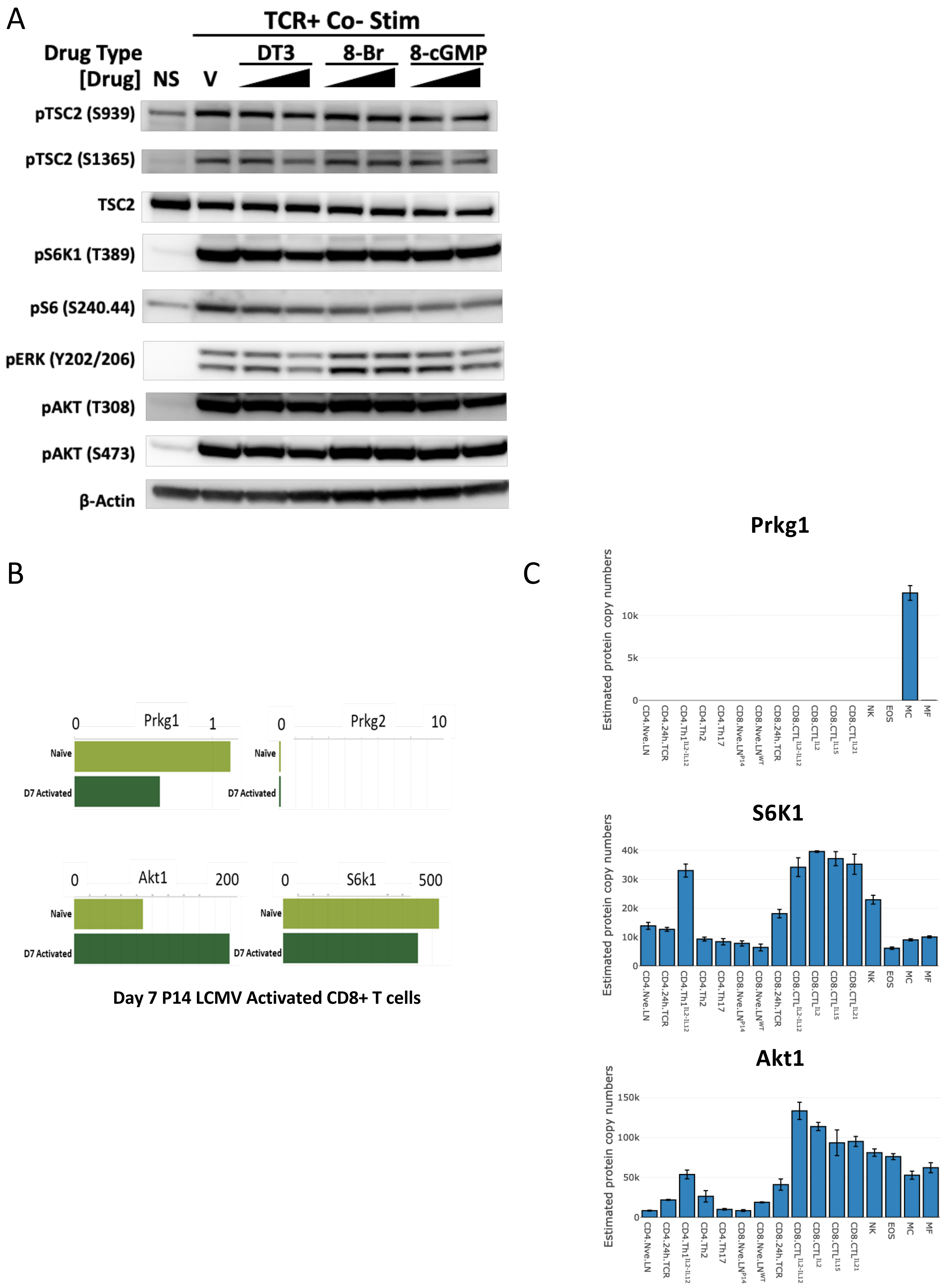
A
TCR+ Co- Stim
Drug Type
[Drug]
NS
V
DT3
8-Br
8-cGMP
pTSC2 (S939)
pTSC2 (S1365)
TSC2
pS6K1 (T389)
pS6 (S240.44)
pERK (Y202/206)
pAKT (T308)
pAKT (S473)
β-Actin
B
Prkg1
0
1
Naïve
D7 Activated
Prkg2
0
10
Naïve
D7 Activated
Akt1
0
200
Naïve
D7 Activated
S6k1
0
500
Naïve
D7 Activated
Day 7 P14 LCMV Activated CD8+ T cells
C
Prkg1
Estimated protein copy numbers
0
5k
10k
S6K1
Estimated protein copy numbers
0
10k
20k
30k
40k
Akt1
Estimated protein copy numbers
0
50k
100k
150k
CD4.Nve.LN
CD4.24h.TCR
CD4.Th1 IL2-IL12
CD4.Th2
CD4.Th17
CD8.Nve.LN P14
CD8.Nve.LN WT
CD8.24h.TCR
CD8.CTL IL2-IL12
CD8.CTL IL2
CD8.CTL IL15
CD8.CTL IL21
NK
EOS
MC
MF

Supplemental Figure 3: cGMP and cGK-1 do not play a role in T cells. A) Immunoblot analysis of T cells stimulated with Signal 1 (TCR, αCD3) and Signal 2 (Co-Stim, αCD28) with increasing concentrations of cGMP agonists. B) Immgen database analysis of Prkg1, Prkg2, Akt1, and S6K1 mRNA expression levels in naïve and D7 activated CD8+ T cells. C) Proteomics analysis of Prkg1, Prkg2, Akt1, and S6K1 mRNA protein levels in various immune subsets Data are representative of 2 experiments (A)

### Sup. Figure 4

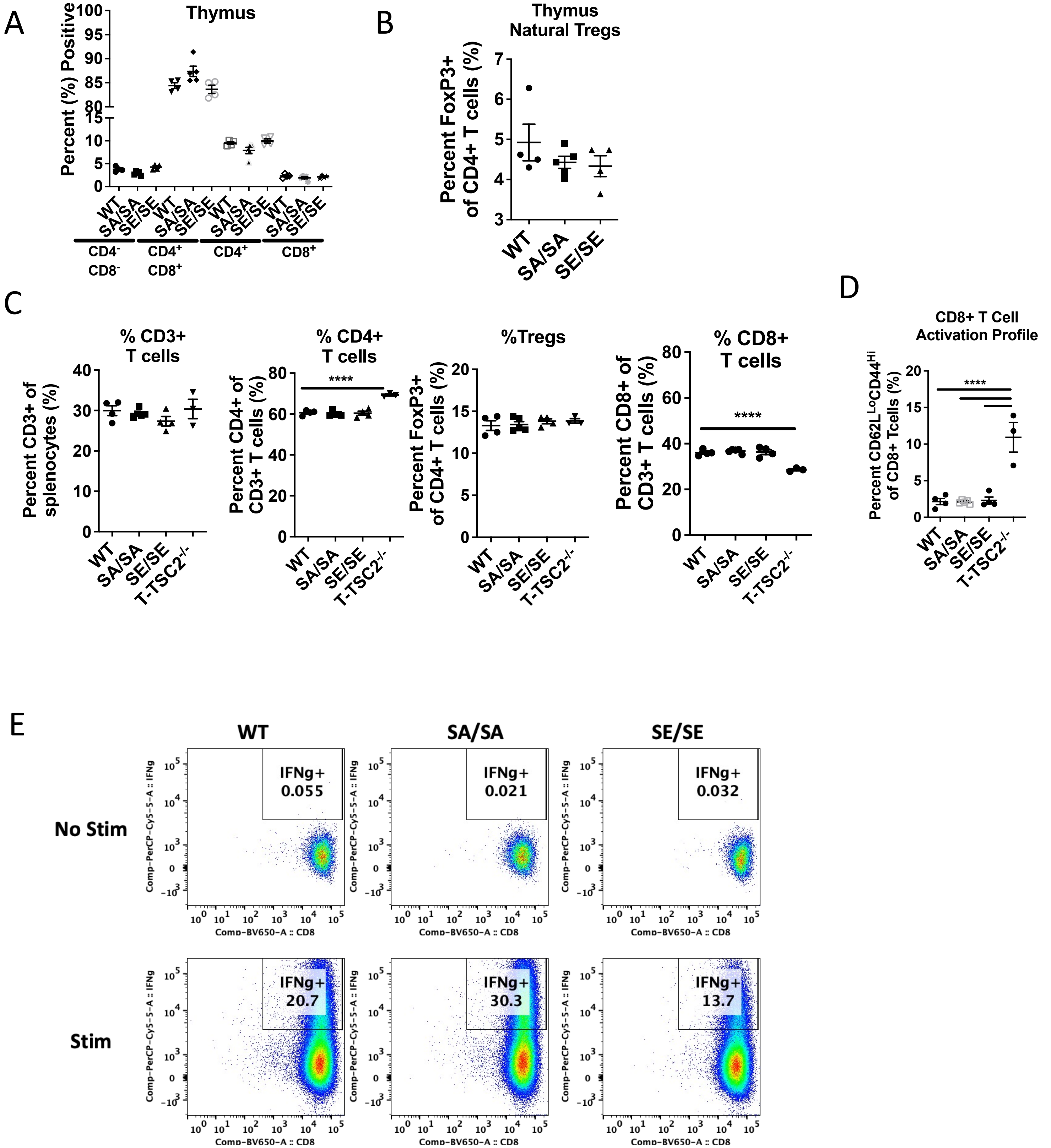
A
Thymus
Percent (%) Positive
WT
SA/SA
SE/SE
CD4- CD8-
CD4+ CD8+
CD4+
CD8+
B
Thymus Natural Tregs
Percent FoxP3+ of CD4+ T cells (%)
C
% CD3+ T cells
Percent CD3+ of splenocytes (%)
% CD4+ T cells
Percent CD4+ of CD3+ T cells (%)
%Tregs
Percent FoxP3+ of CD4+ T cells (%)
% CD8+ T cells
Percent CD8+ of CD3+ T cells (%)
T-TSC2-/-
****
D
CD8+ T Cell Activation Profile
Percent CD62L^Lo CD44^Hi of CD8+ Tcells (%)
E
WT
SA/SA
SE/SE
No Stim
Stim
IFNg+ 0.055
IFNg+ 0.021
IFNg+ 0.032
IFNg+ 20.7
IFNg+ 30.3
IFNg+ 13.7
Comp-PerCP-Cy5-5-A :: IFNg
Comp-BV650-A :: CD8

Supplemental Figure 4: Characterization of lymphocyte development and maturation of age and sex matched WT, TSC2 (SA/SA), TSC (SE/SE) and T-TSC2-/-(TSC2 KO) mice. A) Characterization of lymphocyte development and maturation of age and sex matched WT, TSC2 (SA/SA), and TSC (SE/SE) mice. Flow cytometric phenotyping data compiled from multiple mice (n=4-5) of double negative (CD4-CD8-), double positive (CD4+CD8+), or single positive (CD4+ or CD8+) thymocytes. B) Flow cytometric phenotyping data of thymic Tregs. C) Flow cytometric phenotyping data compiled from multiple mice (n=3-5) of percent CD3+, CD4+, Foxp3+ (CD3+CD4+), CD8+ lymphocytes in the spleen. D) Flow cytometric phenotyping data of activation profile defined by CD62LLoCD44Hion CD3+CD8+T cells in spleen. E) WT, SA/SA, and SE/SE CD8+ T cells were activated and expanded in IL-2 to generate effector cytotoxic lymphocytes for cytokine analysis of IFN-γ upon re-stimulation. Data are representative of one experiment (A-D), two independent experiments (E).

### Sup. Figure 5

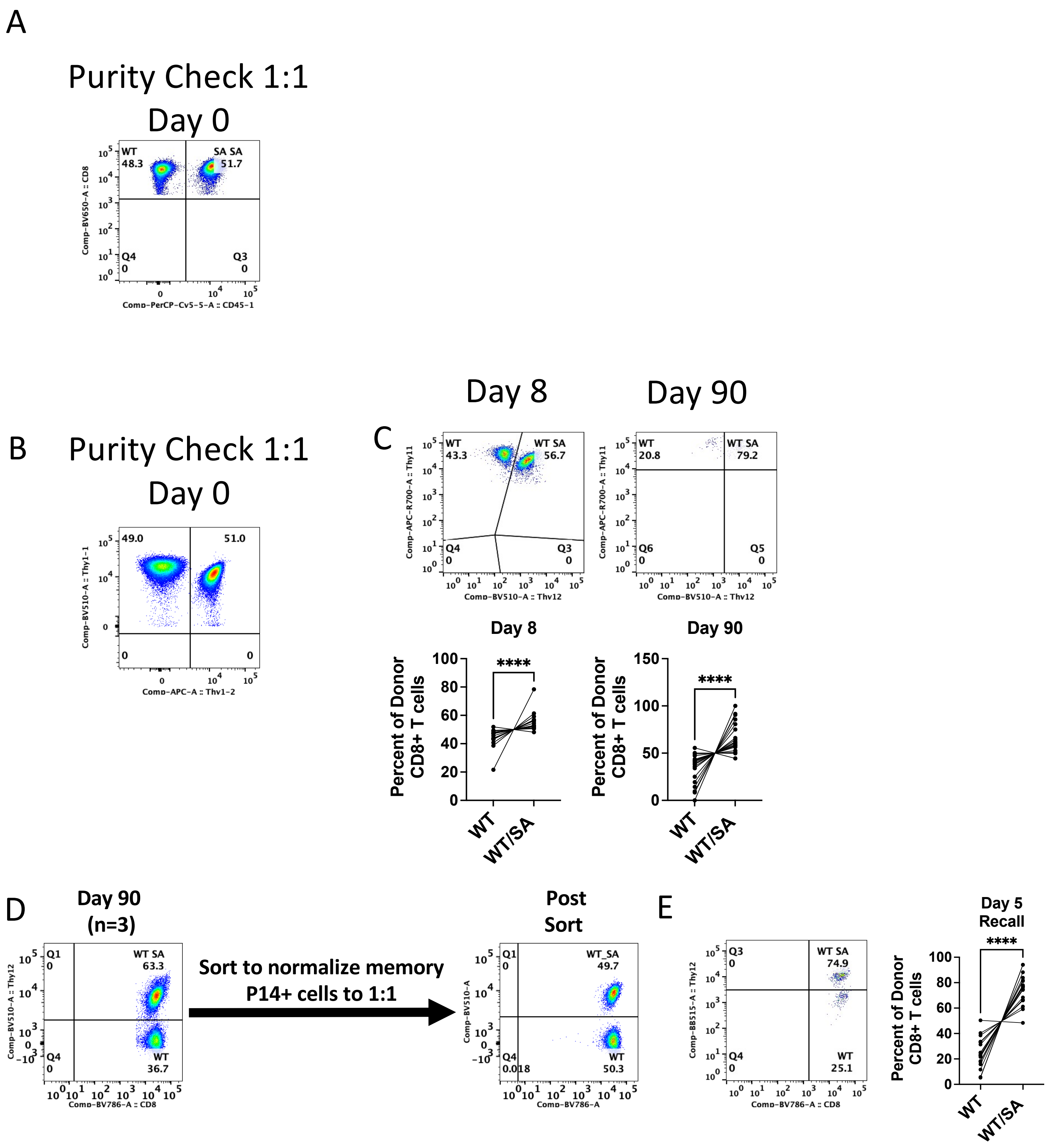

A
Purity Check 1:1
Day 0
WT
48.3
SA SA
51.7
Q4
0
Q3
0
Comp-BV650-A :: CD8
Comp-PerCP-Cy5-5-A :: CD45-1
B
Purity Check 1:1
Day 0
49.0
51.0
0
0
Comp-BV510-A :: Thy1-1
Comp-APC-A :: Thy1-2
C
Day 8
Day 90
WT
43.3
WT SA
56.7
Q4
0
Q3
0
Comp-APC-R700-A :: Thy11
Comp-BV510-A :: Thy12
WT
20.8
WT SA
79.2
Q6
0
Q5
0
Day 8
Day 90
Percent of Donor CD8+ T cells
****
WT
WT/SA
D
Day 90
(n=3)
Q1
0
WT SA
63.3
Q4
0
WT
36.7
Comp-BV510-A :: Thy12
Comp-BV786-A :: CD8
Sort to normalize memory
P14+ cells to 1:1
Post
Sort
Q1
0
WT_SA
49.7
Q4
0.018
WT
50.3
Comp-BV510-A
Comp-BV786-A
E
Q3
0
WT SA
74.9
Q4
0
WT
25.1
Comp-BB515-A :: Thy12
Comp-BV786-A :: CD8
Day 5
Recall
****
Percent of Donor CD8+ T cells
WT
WT/SA

Supplemental Figure 5: Mutating TSC2 at S1365 (SA) promotes strong CD8+ T cells. A) As in Figure 4A, pre-transfer check of co-adoptive WT and TSC2 SA/SA donor OTI CD8+ T cells. B-D) WT and mutant TSC2 (SA) transgenic CD8+ T cells were co-adoptively transferred into naïve WT hosts followed with acute pathogen infection. B) Pre-transfer check of co-adoptive WT and TSC2 WT/SA donor OTI CD8+ T cells C) Flow cytometry plot of transferred CD8+ T cells (top) and summary data (bottom) showing percent of WT vs WT/SA genotype from donor population 8 and 90 days after exposure to Listeria-OVA. D) Percent of memory WT and mutant TSC2 WT/SA P14 CD8+ T cells in spleen (left, n=3). Equal number of memory donor CD8+ T cells were sorted and again co-adoptively transferred into naïve WT recipients to assess memory recall ability on 1:1 basis upon LCMV Armstrong infection. E) Percent of donor CD8+T cells after Day 5 upon recall. ****$p<0.0001$. A paired T-test was performed for statistical analysis. Data are representative of at least 2 independent experiments.

### Sup. Figure 6

A

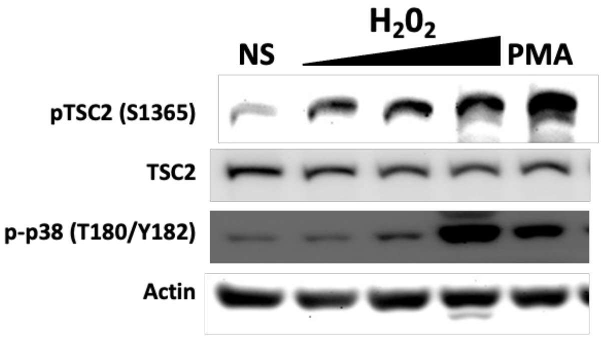


B

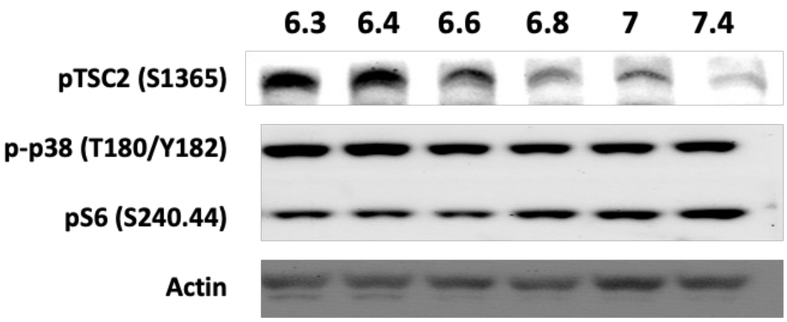

Supplemental Figure 6: Cellular stress activates pTSC2 (S1365) in human T cells. A) Human T cells were exposed to increasing concentrations (.01mM, .1mM, 1mM) of $H_2O_2$ for 30 minutes and assayed by immunoblot analysis as indicated. PMA for 30 minutes is a positive control. B) Activated normal human T cells were exposed to various pH level media conditions for 90 minutes and assayed by immunoblot analysis for pTSC2 and mTORC1 activity. Data are representative of 2 independent experiments.

**Sup. Figure 7**

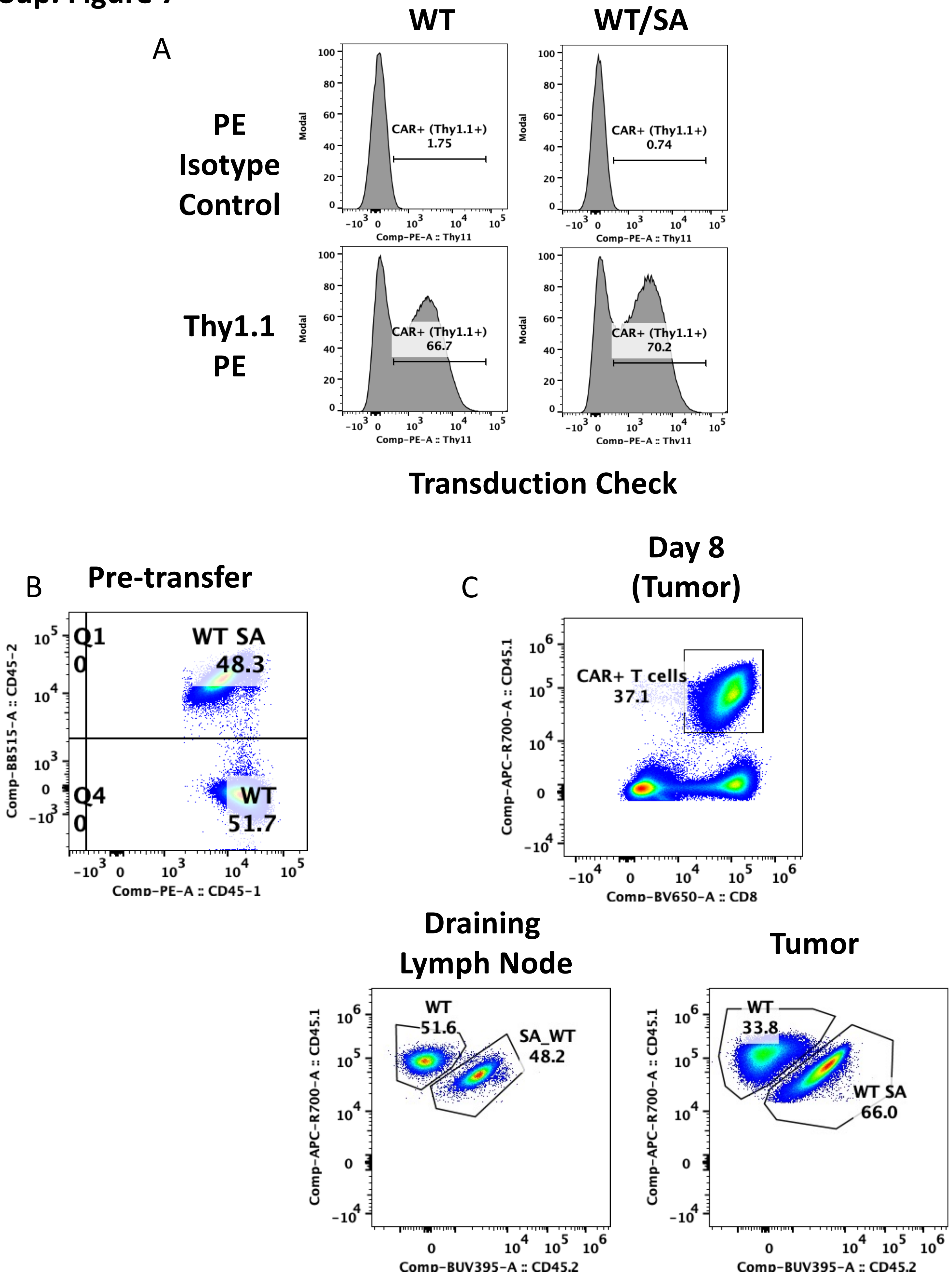
A
WT
WT/SA
PE Isotype Control
Thy1.1 PE
CAR+ (Thy1.1+)
1.75
0.74
66.7
70.2
Modal
Comp-PE-A :: Thy11
Transduction Check
B
Pre-transfer
Q1
0
WT SA
48.3
Q4
0
WT
51.7
Comp-BB515-A :: CD45-2
Comp-PE-A :: CD45-1
C
Day 8 (Tumor)
CAR+ T cells
37.1
Comp-APC-R700-A :: CD45.1
Comp-BV650-A :: CD8
Draining Lymph Node
WT
51.6
SA_WT
48.2
Tumor
WT
33.8
WT SA
66.0
Comp-BUV395-A :: CD45.2

Supplemental Figure 7: SA mutation improves murine CD19 CAR T cells for adoptive cell therapy. A) Transduction efficiency of CAR surface expression assessed by Isotype or Thy1.1 expression via flow cytometry in Figure 6 between WT and TSC2 mutant CD8+ T cells. B) As shown in Figure 6D, pre-transfer efficiency check before transfer into tumor bearing mice. C) As in Figure 6D, representative flow plot of donor TILs in B16-CD19 tumors (Day 8). Data are representative of at least 2 independent experiments.

### Sup. Figure 8

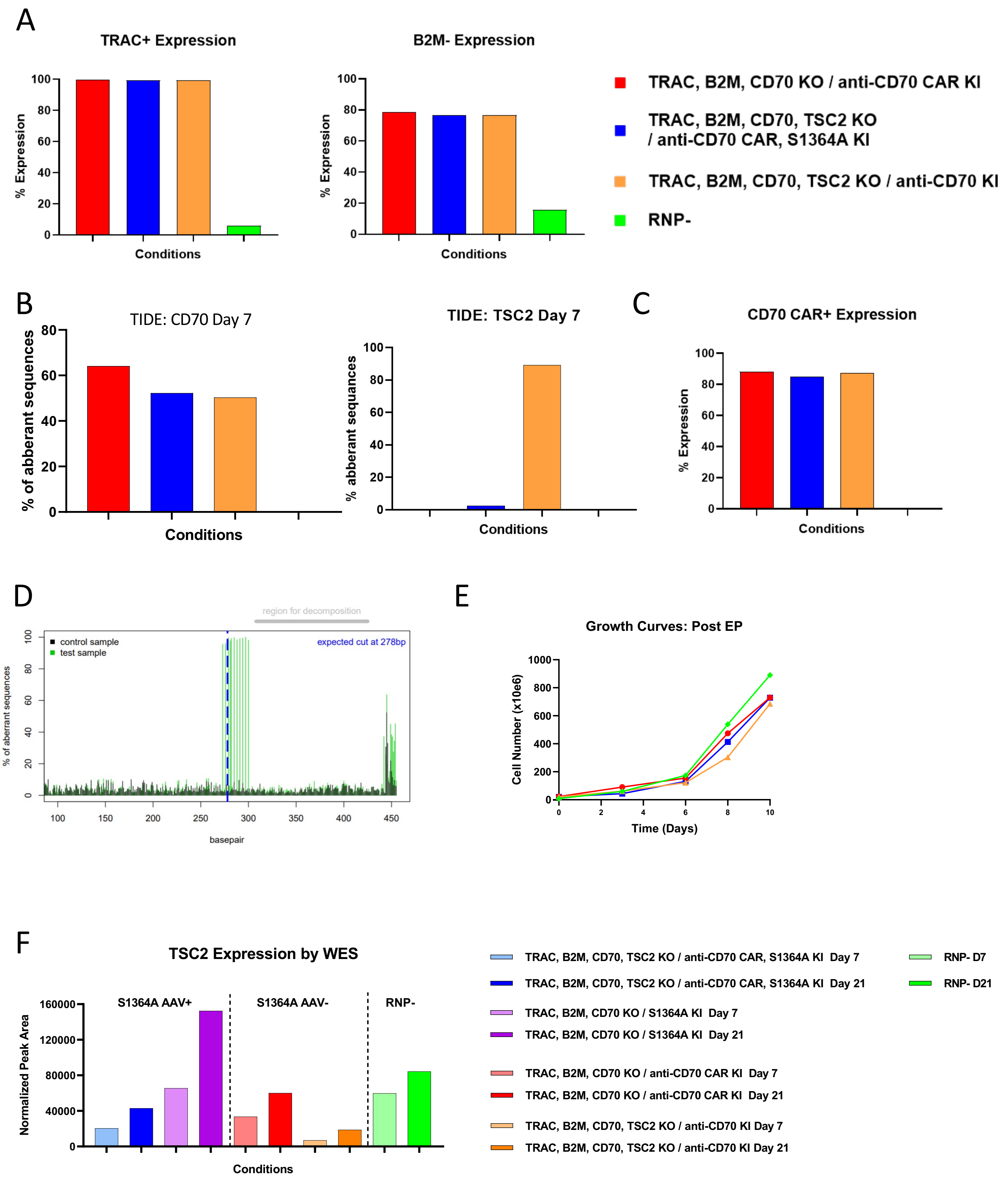
A
TRAC+ Expression
% Expression
Conditions
B2M- Expression
TRAC, B2M, CD70 KO / anti-CD70 CAR KI
TRAC, B2M, CD70, TSC2 KO / anti-CD70 CAR, S1364A KI
TRAC, B2M, CD70, TSC2 KO / anti-CD70 KI
RNP-
B
TIDE: CD70 Day 7
% of abberant sequences
TIDE: TSC2 Day 7
C
CD70 CAR+ Expression
D
region for decomposition
control sample
test sample
expected cut at 278bp
% of aberrant sequences
basepair
E
Growth Curves: Post EP
Cell Number (x10e6)
Time (Days)
F
TSC2 Expression by WES
Normalized Peak Area
S1364A AAV+
S1364A AAV-
RNP-
Conditions
TRAC, B2M, CD70, TSC2 KO / anti-CD70 CAR, S1364A KI Day 7
TRAC, B2M, CD70, TSC2 KO / anti-CD70 CAR, S1364A KI Day 21
TRAC, B2M, CD70 KO / S1364A KI Day 7
TRAC, B2M, CD70 KO / S1364A KI Day 21
TRAC, B2M, CD70 KO / anti-CD70 CAR KI Day 7
TRAC, B2M, CD70 KO / anti-CD70 CAR KI Day 21
TRAC, B2M, CD70, TSC2 KO / anti-CD70 KI Day 7
TRAC, B2M, CD70, TSC2 KO / anti-CD70 KI Day 21
RNP- D7
RNP- D21

Supplemental Figure 8: Engineering CD70 CAR T cells with TSC2 (S1364A) mutation. A) TRAC and B2M knockout was verified by FACS. In cells in which the TRAC loci were edited, the cell populations showed TRAC knock out in less 99.6% of cells, and 79% B2M knock-out surface expression by flow cytometry. B) TIDE analysis verification of CD70 and TSC2 disruption by Tide analysis. C) CD70 CAR expression.  D) TIDE analysis with S1363A donor template (~100% detection of mutation). E) Growth kinetics of CD70 CAR-T cells that were expanded for 10 days. F) Western blot analysis to confirm TSC2 deletion. Data was performed one time (A-F)

### Sup. Figure 9

A

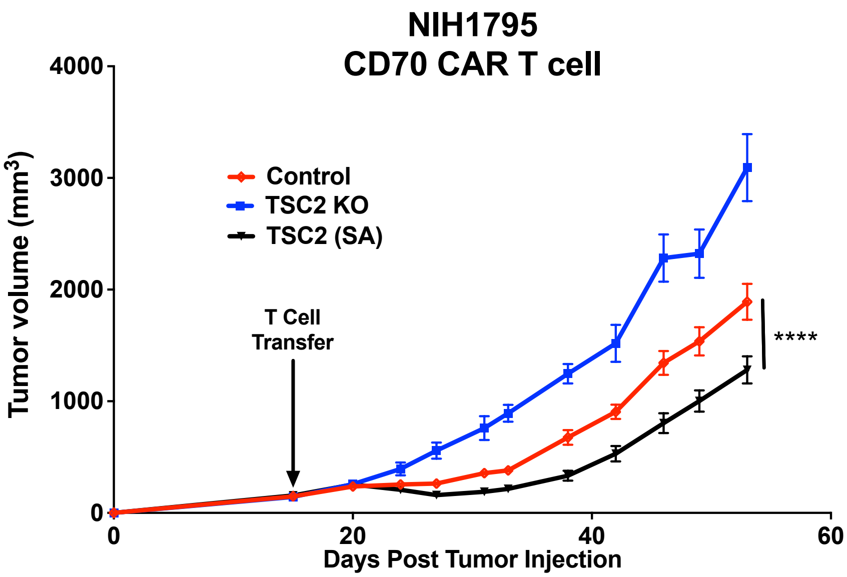


B

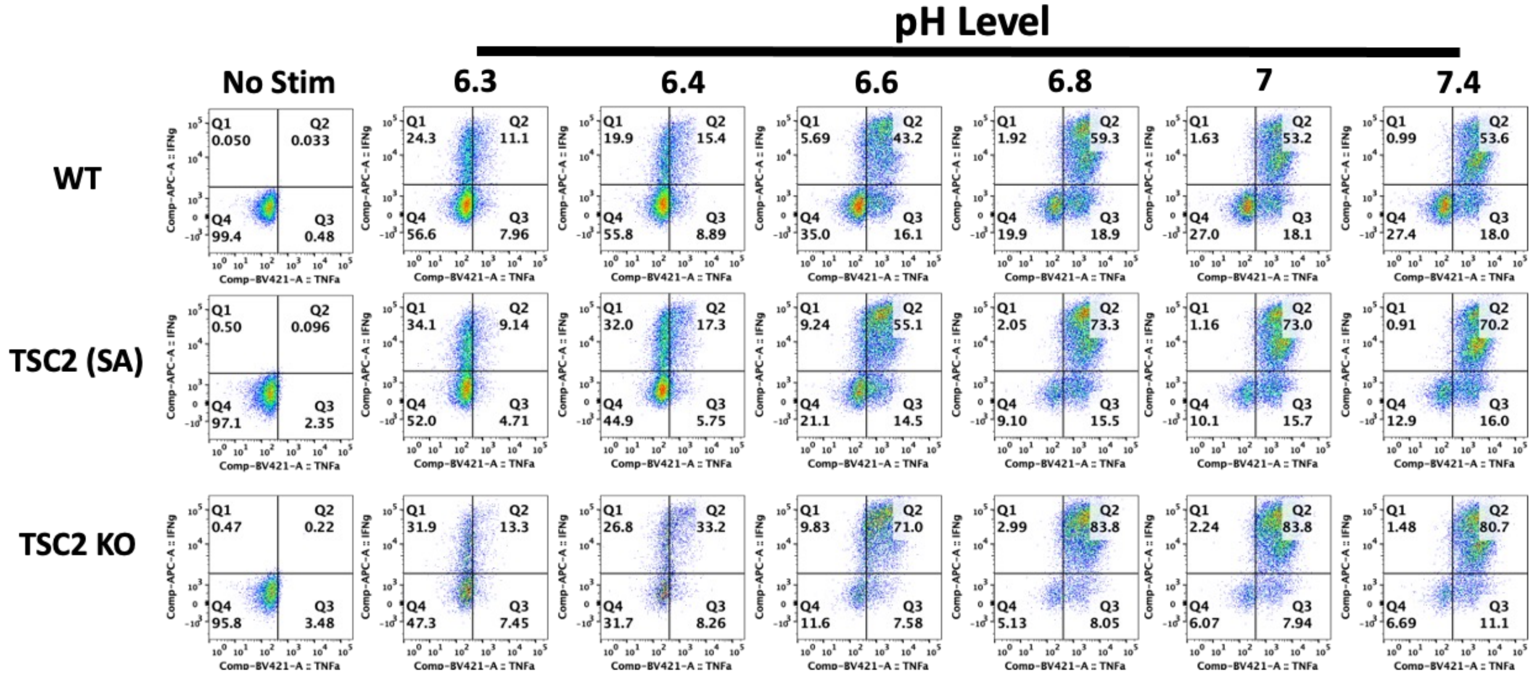


C

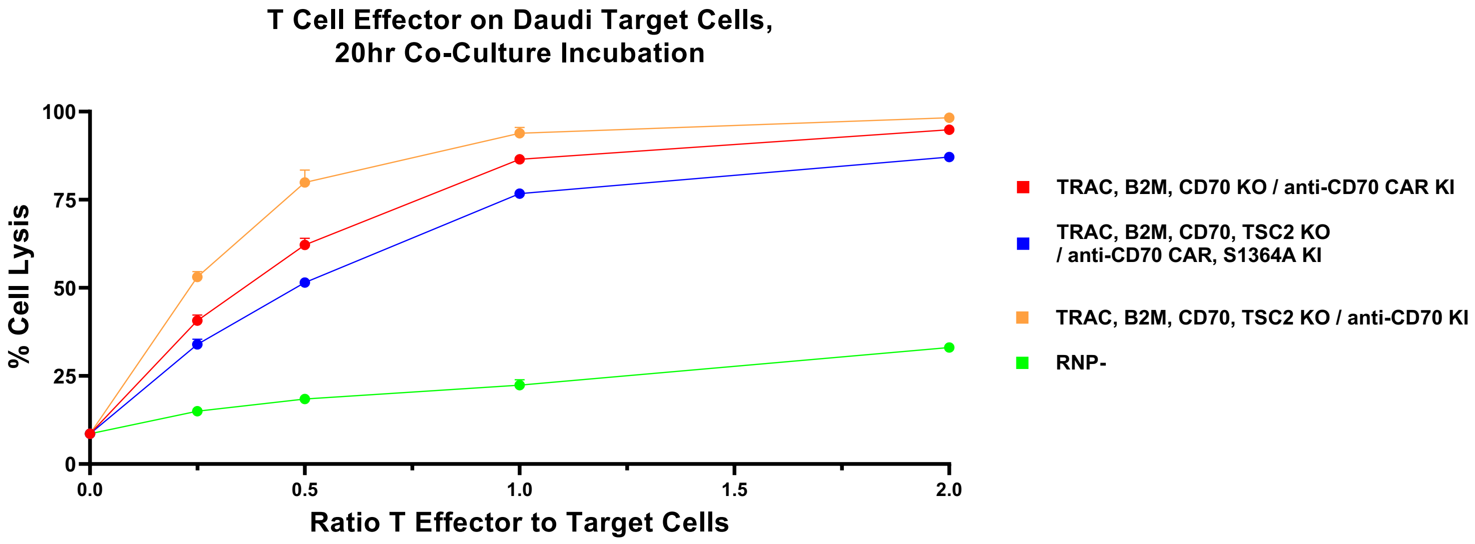

Supplemental Figure 9: SA mutation improves human CD70 CAR T cells to tumor. A) As in 6F but using less donor human CD70 CAR T cells towards CD70 expressing human tumor in NSG mice. B) Control TSC2 SA/SA, or $TSC^{-/-}$ CD8+ T cells were stimulated with PMA and Ionomycin in various pH level media to assess IFN-γ and TNFα via flow cytometry. C) In vitro CTL co-culture using Daudi target cell at a range of CAR T cells to target cell ratios. ****$p<0.0001$ Data was performed one time (A, C) and two independent times (B)
